# Transcriptional readthrough precedes alternative splicing programs triggered in CML cells by imatinib

**DOI:** 10.1101/2025.11.04.686193

**Authors:** Paulina Podszywałow-Bartnicka, Morgan Shine, Jing Lin, Karla M. Neugebauer

**Author notes:** Co-correspondence.

## Abstract

Cellular stresses induce transcription readthrough, whereby RNA polymerase II elongates past a gene’s polyadenylation cleavage site without RNA cleavage. Readthrough has been reported in several cancer types. Here, we use long-read sequencing of nascent RNA to quantify transcriptional readthrough in chronic myelogenous leukemia (CML) cells and characterize early responses to the targeted therapeutic, imatinib. We show that the amount, length, and gene-specificity of readthrough increase within 1 hour, while gene expression and alternative splicing alterations emerge later. Strikingly, imatinib-dependent mRNA isoform changes involved “readthrough chimeras”, in which exons from an upstream gene are alternatively spliced to exons in a downstream gene. Modifications in mRNA isoforms and chimera levels detected at 18 hours were also found in imatinib-resistant K562 as well as CML patient cells, suggesting a cascade of early changes in the fidelity of transcription and splicing, leading to long-term adjustment in gene expression and the development of therapy resistance.

**Teaser:** Precision RNA sequencing was used to discover the earliest response of leukemia cells treated with targeted therapy: transcriptional readthrough.

## INTRODUCTION

From prokaryotes to eukaryotes and from yeast to man, nascent RNA is the substrate of numerous RNA processing steps, including 5’ end capping, splicing, and 3’end cleavage [1]. Specific RNA sequences recruit the corresponding machineries – namely the capping enzymes, the spliceosome, and the Cleavage and Polyadenylation Complex (CPC) – that act quickly to modify the nascent transcript as it emerges from RNA polymerase II (Pol II). Human genes harbor an average of 8 introns each, of which approximately 75% are co-transcriptionally removed by the spliceosome shortly after the synthesis of 5’ and 3’ splice sites delineating each intron [1, 2]. The recent introduction of long-read sequencing of nascent RNA as a tool for analyzing co-transcriptional RNA processing has enabled researchers to associate multiple RNA processing steps with the position of Pol II. Emerging evidence points to tight coordination of intron removal at the level of individual transcripts [3–6]. Recently, “all-or-none” transcripts were described in budding and fission yeasts as well as mouse erythroblasts, in which the retention of multiple introns in the same mRNA molecule is associated with transcriptional readthrough, in which Pol II proceeds past the polyadenylation cleavage site (PAS) and continues downstream of the parent gene [4, 7, 8]. A simple interpretation of this phenotype is that the activity of CPC is either delayed or blocked when a transcript has not been co-transcriptionally spliced.

Transcriptional readthrough has emerged as a common, rapid response to cellular stress. Hypoxia, heat shock, starvation, and viral infection each lead to increases in the level of intergenic transcription, as measured by RNA sequencing experiments that identify “downstream of genes” (DoG) RNAs [9, 10]. Increased intergenic transcription can encompass many kilobases (kb) past the most distal gene boundary and is believed to be contiguous with the upstream “parent” gene rather than activation of new transcription start sites in intergenic regions [11, 12]. When transcription continues into the next gene on the DNA strand, transcription and/or splicing of the “read-in gene” can be disrupted [8, 13, 14]. Although DoG RNAs were considered to be stress-induced, their presence has since been broadly documented in normal tissues, cancer cells, and aging cells and tissues [15–17]. Broadly speaking, the cellular role(s) of DoG RNAs is unknown. However, transcriptional readthrough is a hallmark of clear cell renal carcinoma cells and prostate cancer cells, where a marked increase in transcription downstream of genes leads to the formation of chimeric RNA transcripts comprised of exons from the parent gene spliced together with exons from the read-in gene [18, 19]. Chimeric transcripts can encode truncated proteins, newly fused together functional globular domains or, alternatively, proteins with intrinsically disordered extensions [17]. Thus, the induced expression of chimeric transcripts and encoded proteins in response to stress may be physiologically relevant.

Pre-mRNA splicing regulation is often disrupted in cancer and even drives cancer initiation in some blood cancers, making splicing a target for clinical therapies [20]. By contrast, transcriptional readthrough has been vastly understudied in the context of cancer. We are interested in the possibility that transcriptional readthrough and potential downstream changes in RNA processing play roles in the development and treatment of chronic myeloid leukemia (CML), where alternative splicing has been identified in patients [21]. Activation of cellular stress responses is detected in stem cells and accompanies carcinogenesis [22, 23]. These evolutionarily conserved pathways are detected in primary cells and hijacked by different cancer types, supporting survival and chemotherapy resistance. In CML, the unfolded protein response and phosphorylation of eIF2α, hallmarks of endoplasmic reticulum stress, accompany cancer progression and reduce the effectiveness of the targeted therapeutic drug imatinib [24, 25]. The activity of Bcr-Abl1 onco-kinase affects protein translation and induces stress granules in CML cells [26, 27]. Similar cellular phenotypes were detected in acute myeloid leukemia (AML), linking condensation of stress granules to changes in splicing regulation [28]. Two chemotherapeutic agents used to treat AML, among other cancers, JTE-607 and camptothecin, cause transcriptional readthrough by directly inhibiting the enzymatic activity of the CPC and topoisomerase 1, respectively [29–32]. Because these chemotherapeutic compounds directly and indirectly impact transcription and pre-mRNA splicing [33–35] it is unclear if transcriptional readthrough is purely related to their biochemical activities or is related to the stress induced in cancer cells by therapeutic treatment.

Here, we have specifically addressed the possibility that transcriptional readthrough is induced by imatinib treatment of CML cells. Because imatinib targets the cytoplasmic kinase Abl1 and *not* the transcription or RNA processing machinery, our experiment instead enables us to determine the transcriptional effects of cancer cells stressed by targeted therapy. To achieve this, we have also implemented long-read sequencing of nascent RNA together with short-read sequencing datasets to quantify and better characterize the occurrence of transcriptional readthrough at the gene level and at the level of single transcripts, enabling us to detect associated events like alternative splicing and changes in the formation of chimeric transcripts. Taken together, our findings show that transcriptional readthrough is the first detectable effect of imatinib on gene expression, which precedes and facilitates changes in alternative splicing.

## RESULTS

### The targeted chemotherapeutic imatinib induces transcriptional readthrough in K562 cells

To study the effects of imatinib on RNA processing, we chose the K562 cell line, which originates from a patient with chronic myeloid leukemia in blast crisis (CML-BC) [36], as a model system. Transcriptional readthrough is typically studied by sequencing nuclear RNA with a short-read (e.g. Illumina) platform [32]. In this study, we implemented long-read sequencing of nascent RNA to detect and quantify the number and length of reads that extend past the PAS relative to the number that have 3’ ends within the gene body (Fig. 1A). We reasoned that levels of transcriptional readthrough might be altered by imatinib treatment compared to control and that the chemical inhibitor of CPC, JTE-607, could serve as a positive control for increased transcriptional readthrough of at least a subset of genes [32, 37]. Nascent RNA long-read libraries were prepared by isolating chromatin fractions (Fig. S1A) from three biological replicates each for four different treatments: control (CTRL), 1-hour treatment with JTE-607 (JTE 1h), 1-hour treatment with imatinib (IM 1h), and 18-hour treatment with imatinib (IM 18h) [38]. Read numbers, mapping statistics, and read lengths are available in Tables S1 and S2; the read length distributions (Fig. S1B), similar to previous nascent RNA long reads, allow us to analyze RNA processing early in gene transcription and additionally capture longer reads that might reflect transcriptional readthrough for approximately 11,0000 genes per dataset (Fig. S1C). Normalization indicates a similar read length distribution and number of representative genes across all datasets (Figs. S1D,E). Meta-analysis of the resulting read coverage across the gene body and extending 4 kb upstream of the transcription start site (TSS) and downstream of the last PAS revealed that JTE-607 treatment led to transcriptional readthrough, as expected (Fig. 1C). Note that the decrease in transcriptional readthrough approaching +4 kb is in part determined by the length limitations of long-read sequencing (Fig. S1F, Fig. S1B, D). Taken together, IM 1h and 18h treatments were characterized by higher levels of readthrough compared to CTRL and lower levels than JTE. These data raise the question of which genes are affected by imatinib.

**Figure 1.**
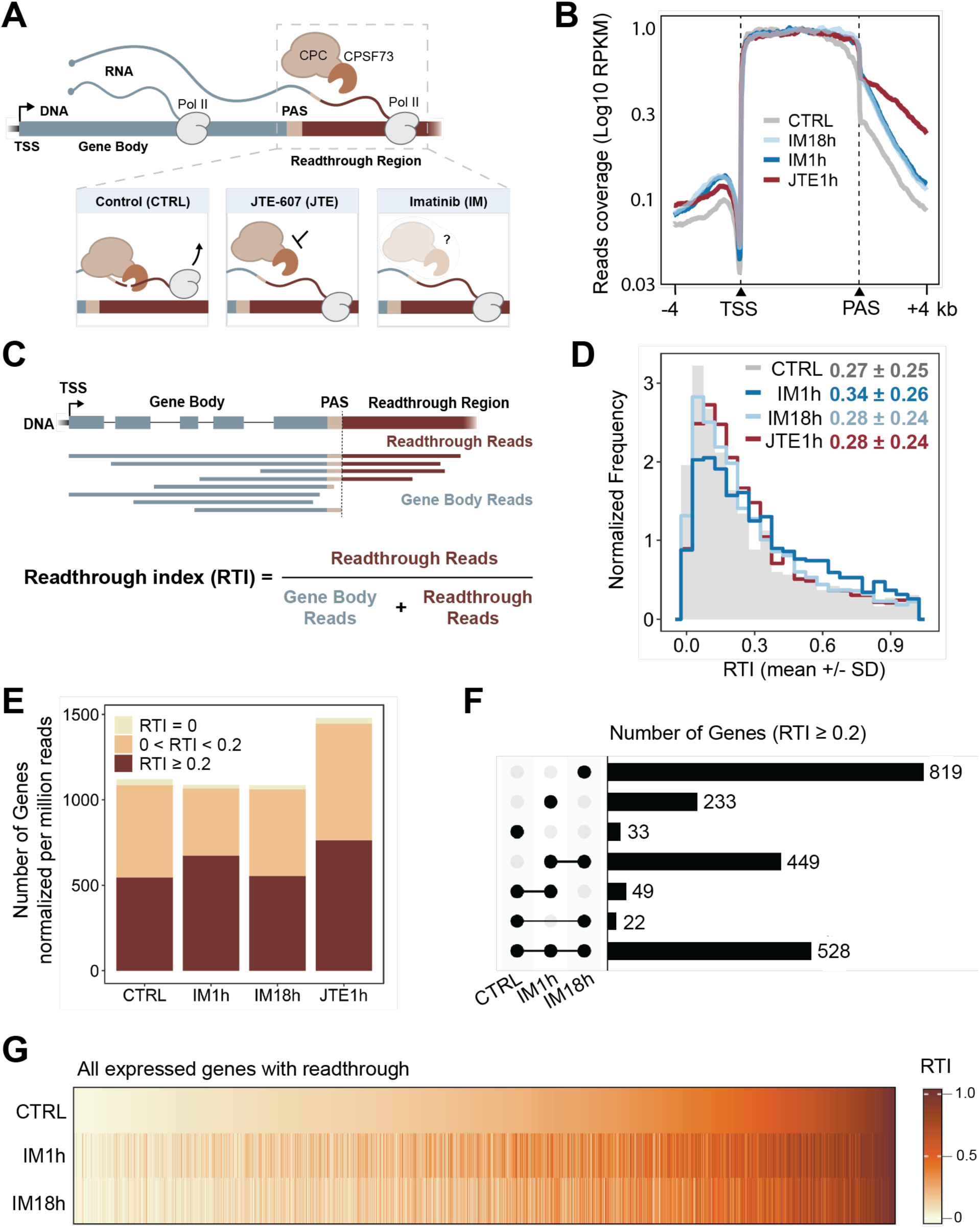
Imatinib treatment induces transcriptional readthrough detected by long-read sequencing. **(A)** Schematic showing transcriptional readthrough, which occurs upon impaired 3’ end cleavage. JTE-607 is a chemical inhibitor of CPSF73, and imatinib is a targeted therapeutic that inhibits the Bcr-Abl1 onco-kinase. PAS - polyadenylation signal, TSS - transcription start site, Pol II - RNA polymerase II. **(B)** Metagene analysis of RNA-Seq profiles for control (CTRL) and treated (IM 1h, IM 18h, JTE 1h) cells. The y-axis represents the read coverage combined from three biological replicates and analyzed in 40-nucleotide bins. RPKM – Reads Per Kilobase per Million mapped reads, TSS - transcription start site, PAS - polyadenylation signal. **(C)** Schematic representation of transcriptional readthrough index (RTI) calculation. **(D)** Distribution of RTI values per gene for each condition. RTI values (mean ± s.d.) were calculated based on 3 biological replicates per condition and are provided in the top right corner of the plot. **(E)** Fraction of genes with more or less than 20% of long reads displaying readthrough (RTI = 0 in beige; RTI < 0.2 in yellow; RTI ≥ 0.2 in maroon). In panels D and E, genes were required to have ≥ 10 reads/replicate to be included in these analyses. **(F)** Upset plot comparing genes with RTI ≥ 0.2 in CTRL, IM 1h, and IM 18h cells. **(G)** Heat map of RTI values for each expressed gene (n=1978 genes from panel **E**), ordered by RTI value in CTRL.

To generate parameters from our data that would enable gene-specific analysis of nascent RNAs sequenced in our libraries, we defined readthrough reads as those that originate at either the TSS or within the gene but have 3’ ends ≥100 nucleotides downstream of the last annotated PAS (Fig. 1C). We defined reads with 3’ ends within the transcribed gene (i.e. between the TSS and the PAS) as “gene body” reads (gb). This allowed us to derive a gene-specific metric, the Readthrough Index (RTI), which signifies the proportion of long reads that are not cleaved efficiently. Comparing across treatment groups (Fig. 1D), genes with RTI>0 had a similar distribution of RTI values, with a trend to higher RTIs in the IM 1h dataset. JTE 1h treatment was characterized by overall more genes (∼1,450/million reads) with RTI>0 compared with CTRL, IM 1h, and IM 18h, which each had 1,000-1,100 genes/million reads with readthrough (Fig. 1E). These data are consistent with the findings of others that JTE inhibits the cleavage of some but not all genes [32, 37] and show that transcriptional readthrough is detected and quantifiable in nascent RNA long-read sequencing libraries.

To address whether imatinib treatment induces readthrough in different sets of genes, we analyzed unique and overlapping genes with RTI≥0.2 (Fig. 1F). 528 genes were common among CTRL and imatinib treatments. The sample with the most (819) unique genes with readthrough is IM 18h, while 449 genes in IM 1h and 18h were shared and not present in CTRL. To determine how the extent of readthrough varies between CTRL, IM 1h, and IM 18h, we created a heatmap with CTRL RTIs sorted from low to high by gene; corresponding RTI values for imatinib treatments were aligned accordingly (Fig. 1G). The data show a range of altered intensities across the genes analyzed for IM 1h and 18h, with many similarities between the two timepoints. Thus, even though IM 18h has the largest number of unique genes with RTI≥0.2 (Fig. 1F), our data indicate a striking induction of transcriptional readthrough within 1 hour of imatinib treatment.

### Intron retention is associated with readthrough and unaffected by imatinib

Long reads also reveal splicing information that can be correlated with other features of each read, such as transcriptional readthrough, within the same RNA molecule [4, 7, 8]. Therefore, we sought to determine if transcripts with readthrough displayed elevated levels of intron retention and, if true, whether imatinib treatment altered this result. To do so, we quantified co-transcriptional splicing efficiency (CoSE) of CTRL, IM 1h, and IM 18h, by aligning our long reads to gene annotations (Fig. 2A, top panel). Relatively few introns showed substantial changes in CoSE in either the positive (more splicing) or negative (intron retention) direction after 1 or 18 h IM treatment (Fig. 2A, bottom panel) or after JTE treatment (Fig. S2A). Only ∼10% of introns were retained in full-length reads that begin at the TSS and have 3’ ends within the gene body, consistent with efficient co-transcriptional splicing and unaffected by IM 1h or 18h (Fig. 2B). As expected, more introns (∼35%) were retained in readthrough reads, and this was similarly unaffected by imatinib. Because of read length limitations, focusing on full-length reads decreases the number of intron-containing readthrough reads we can analyze. Therefore, we separately analyzed readthrough reads that did not start at the TSS and found ∼40% and ∼75% retained introns in gene body and readthrough categories, respectively. Again, neither parameter was significantly affected by imatinib (Fig. 2B). A partial correlation between transcriptional readthrough and gene-specific CoSE was reinforced by global analysis, reflecting the increased intron retention present in readthrough transcripts (Fig. S2B). We conclude that although readthrough transcripts are correlated with intron retention in human K562 cells, imatinib does not trigger intron retention.

**Figure 2.**
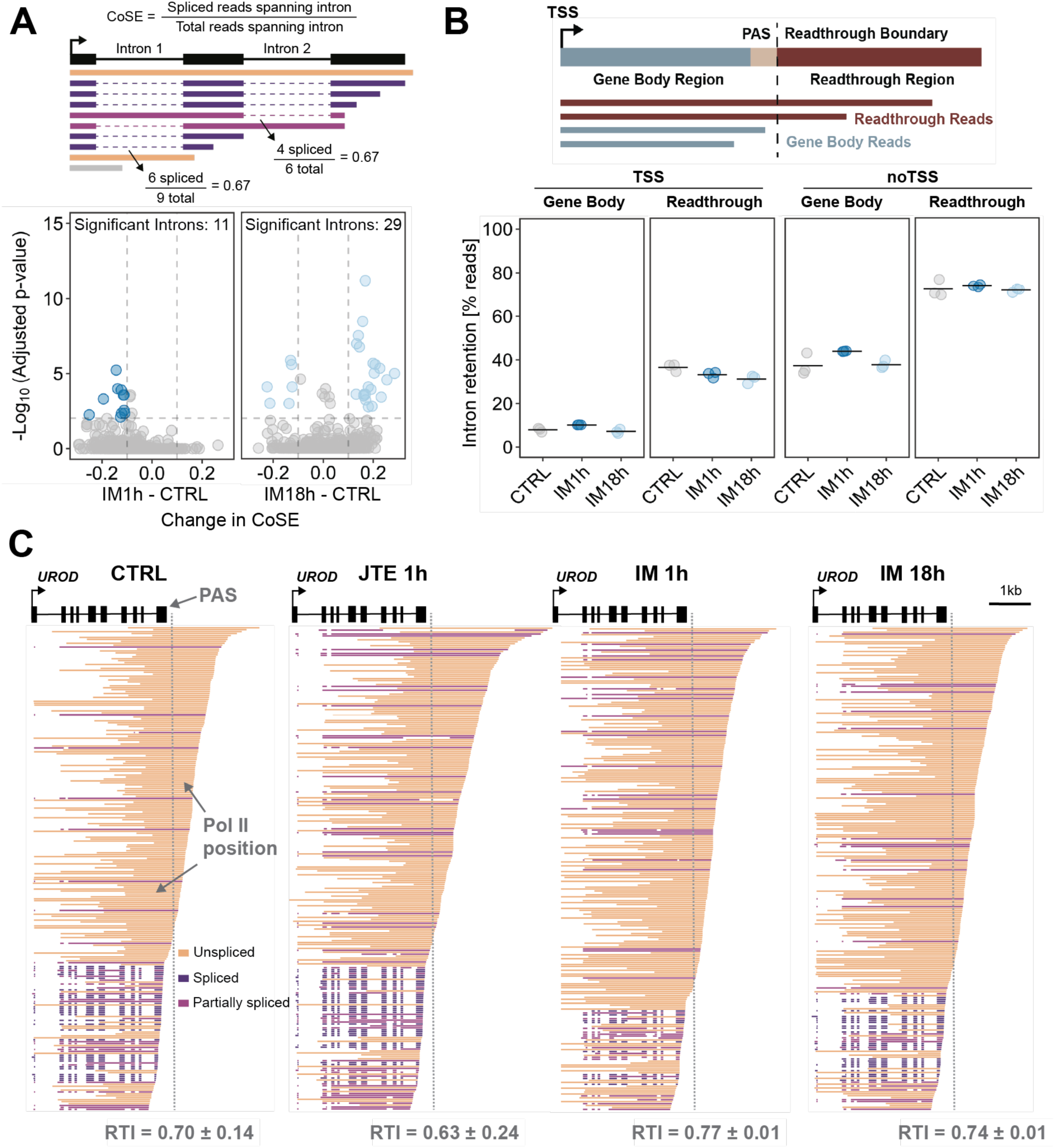
Transcriptional readthrough is associated with higher incidence of intron retention. K562 cells were untreated (CTRL) or treated with Imatinib (IM) or JTE-607 (JTE) for 1h or 18h. **(A)** Analysis of co-transcriptional splicing efficiency (CoSE) based on long-read sequencing of nascent RNA. *Upper panel*: Schematic explanation of CoSE metric. *Lower panel:* Change in CoSE value for IM 1h or 18h IM relative to CTRL plotted for individual introns (n=4662 total introns for 1h IM; 4846 for 18h IM). Introns with significant changes in splicing efficiency (|Change in CoSE| ≥ 0.1 and adjusted p-value < 0.05) are marked in blue. An adjusted p-value was determined for each intron using chi-square tests for pairs of replicates and the Benjamini-Hochberg procedure. **(B)** *Upper panel*: schematic explanation of classification of long reads as “gene body” or “readthrough”. *Lower panel:* Percent nascent RNA long reads with all introns retained (“unspliced”). Reads were either filtered for those beginning within 100 bp of the (TSS) or not filtered (noTSS). The black line represents the mean of 3 biological replicates. **(C)** Alignment of nascent RNA long reads to the gene uroporphyrinogen decarboxylase (*UROD*). Reads are colored according to their splicing status and downsampled to 250 reads total. RTI values (mean ± S.E.M) for *UROD* from three independent experiments are shown. The gray dotted line represents the beginning of the readthrough region, and the 3′ end of each read corresponds to Pol II position. PAS – polyadenylation signal.

In a previous study, we found that transcripts from the β-globin gene and other erythroid genes displayed “all-or-none” RNA processing, in which half of the transcripts were efficiently spliced and cleaved at the 3’ end [4]. The other half contained all of their introns and displayed transcriptional readthrough. We searched for genes displaying “all-or-none” processing and found that UROD exhibits this behavior (Fig. 2C). Alignment of subsampled UROD transcripts show partially spliced, all spliced, and all unspliced transcripts aligned according to the position of Pol II. As expected, the majority of readthrough transcripts are unspliced, and both IM 1h and 18h treatments increase the RTI. The UROD gene encodes the enzyme uroporphyrinogen decarboxylase, which is crucial for heme production. Interestingly, GO term analysis for the genes with RTI≥0.2 in IM 1h and 18h highlights metal ion binding and enzymatic activities characteristic of erythropoiesis (Fig. S3A-B), suggesting that imatinib is able to direct K562 cells into the erythroid lineage on shorter timescales than previously appreciated [39].

### Erythroid differentiation accompanies transcriptional readthrough upon imatinib treatment

To determine the cellular significance of these early gene expression changes in response to imatinib, we generated datasets from short-read polyA+ RNA-seq libraries to corroborate and extend our findings (Table S1). To ask if the genes with high RTI (see Fig. 1G) belonged to previously or newly expressed genes, we analyzed 1818 genes with RTI≥0.2 in IM 18h samples; RTI values per gene were sorted according to the IM 1h dataset to visualize how readthrough and gene expression change over time (Fig. 3A, upper panel). The same 1818 genes were sorted in the same order and then expressed in terms of fold change in mRNA level compared to control, showing that only minor changes in mRNA levels were detected at 1h (Fig. 3A, lower panel). Strikingly, the data at 6h and 18h clearly indicate that 78% of the genes with high RTI values at 1h are downregulated over the time period. In contrast, the genes with low RTI at 1h are mostly upregulated and increase in RTI by 18h, accounting for the increased numbers of readthrough genes by the later timepoint. We conclude that genes with significant readthrough at 1h of imatinib treatment are on a trajectory of changing gene expression.

**Figure 3.**
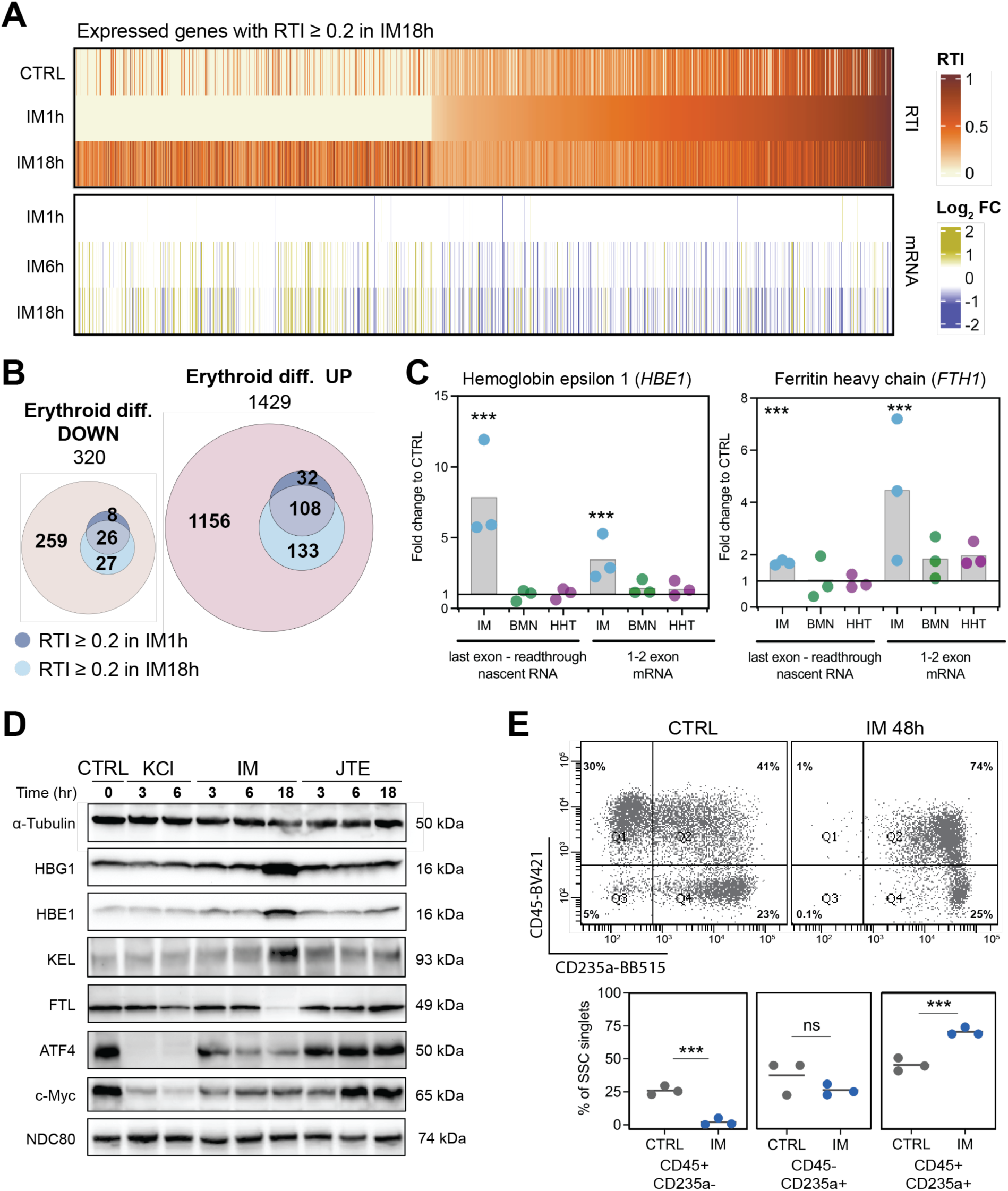
Imatinib induced transcriptional readthrough is present in genes associated with erythropoiesis. **(A)** Changes in RTI and mRNA levels upon IM treatment for genes with RTI ≥ 0.2 in IM18h (n=1818), ordered by RTI in IM1h. Log2FC - log2 of fold change relative to CTRL. **(B)** Prevalence of genes with RTI ≥ 0.2 after IM 1h or 18h overlapping with genes that are up- or down-regulated upon progenitor differentiation into poly- and ortho-erythroid cells (Erythroid diff. UP or DOWN, respectively). Values represent numbers of genes. **(C)** Changes in transcriptional readthrough and mRNA levels upon 18-hour treatment with Imatinib (IM), Talazoparib (BMN), or Homoharringtonine (HHT) quantified for hemoglobin e (*HBE1*) and ferritin heavy chain (*FTH1*) using RT-qPCR. Significance based on two-tailed Student’s t-test (*** p < 0.005). **(D)** Western blot detection of proteins encoded by selected up- and down-regulated genes important in erythroid differentiation and cell cycle regulation. Equal amounts of protein from whole cell lysates were loaded in each lane). Loading control, a-Tubulin. **(E)** Erythroid differentiation markers analyzed by flow cytometry in CTRL and cells treated with IM for 48 hours (IM 48h). Antibodies conjugated to fluorophores BV421 and BB515 were used for detection of CD45 and CD235a proteins, respectively. (**Upper panel**) Representative plot showing percent cells positive for each CD marker; (**lower panel**) black line represents mean value of three replicates (points). Significance based on two-tailed Student’s t-test (*** p < 0.005).

Analysis of differential gene expression revealed an increase in some erythroid genes, such as hemoglobins, even after 1h of imatinib treatment (Fig. S4A). To determine if genes with transcriptional readthrough identified upon IM 1h or 18h in K562 cells belong to the constellation of genes (erythroid or non-erythroid) that are up- or down-regulated during normal erythropoiesis, we utilized a dataset from human CD34^+^ hematopoietic stem cell progenitors (HSPCs) undergoing differentiation into poly- and ortho-erythroid cells [40]. 334 of the identified genes with RTI≥0.2 (16%) overlapped with the normal erythropoiesis dataset, increasing in number from 1h to 18h of imatinib (Fig. 3B). Mean RTI for these overlapping genes reveals a trend to higher RTI in IM 1h (Fig. S4B). We used RT-qPCR to validate two examples – hemoglobin ε (*HBE1*) and ferritin heavy chain (*FTH1*) – that displayed increased transcriptional readthrough and mRNA levels in IM 18h (Fig. 3C). Interestingly, neither of two other chemotherapeutics with distinct mechanisms of action – Talazoparib (BMN) or Homoharringtonine (HHT) – induced transcriptional readthrough, indicating the specificity of imatinib’s effects on 3’ end cleavage, at least for these two genes. To further validate the specific, early induction of erythropoietic genes by imatinib, we performed western blotting and detected elevated globins γ and ε as well as Kell endopeptidase (*KEL*) in IM 18h samples (Fig. 3D). Finally, flow cytometry analysis of K562 cells treated with imatinib for 48h confirmed their differentiation to CD45+ CD235a+ erythroblasts (Fig. 3E), validating and extending previous findings [39].

### Imatinib leads to alternative mRNA isoforms with and without transcriptional readthrough

It is well known that alternative splicing and intron retention are regulatory features of erythropoiesis [40–42]. Since readthrough is an early response to imatinib treatment, we wondered if and when alternative splice isoforms might be detected. To do so, we generated short-read sequencing datasets from total nuclear RNA (Table S3); our rationale was (*i*) changes in splicing will first be detected in the nucleus before achieving steady-state mRNA representation in the cytoplasm and (*ii*) intron retention prevents mRNA export from the nucleus, enabling better detection of this regulatory event in the nuclear fraction. Using splicing per intron (SPI) as a statistically powerful metric, we detected a few intron retention events in IM 1h and a large increase in splicing efficiency at 18h (Fig. 4A), in agreement with the long-read nascent RNA sequencing results (see Fig. 2A). Importantly, many changes in introns spliced out occurred at IM 18h, indicating likely expression of mRNA isoforms produced by alternative splicing. Therefore, we analyzed our mRNA-seq dataset for alternative splicing changes comparing CTRL to IM and JTE 18h using rMATS. This revealed ∼500 alternative splicing events in IM 18h, greater than 10 times the number of changes in IM 1h (Fig. S5A), including a large number of cassette exon events, both inclusion and exclusion (Fig. 4B). As a control for possible direct or indirect effects of transcriptional readthrough on alternative splicing, a dataset was collected for JTE 1h and 18h; approximately 40 and 150 alternative splicing changes were detected, respectively (Fig. S5A). By contrast, another stress condition associated with transcriptional readthrough – hyperosmotic shock – led to almost 400 alternative splicing events, including many cassette exons and retained introns, after only 1h of 110 mM KCl treatment (Fig. S5A-B). We conclude that direct inhibition of 3’ end cleavage by JTE itself leads to significant alternative splicing, suggesting that some of the alternative mRNA isoforms induced by IM could be due to readthrough. In addition, alternative splicing appears to be a robust and specific response to imatinib treatment that differs from JTE treatment and hyperosmotic shock.

**Figure 4.**
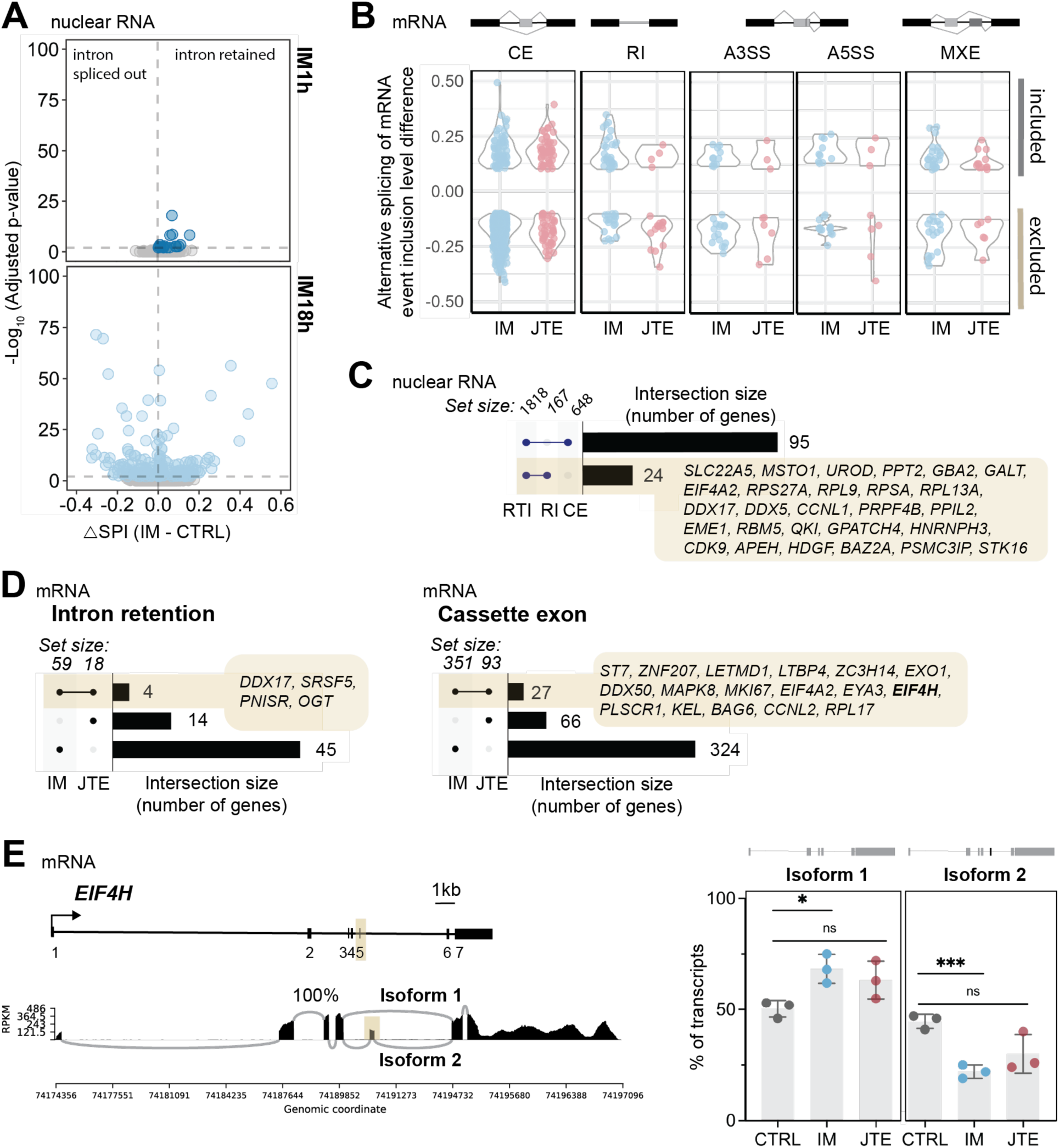
Widespread alternative splicing in response to imatinib correlates with readthrough in some cases. **(A)** Change in SPI values from short-read sequencing of mRNA in IM 1h (upper) or 18h (lower) relative to CTRL. Introns with significant changes in SPI (adjusted p-value < 0.05) are marked in blue. Adjusted p-values were determined using chi-square tests for pairs of replicates and the Benjamini-Hochberg procedure. **(B)** Significant changes in alternative splicing in cells treated with JTE or IM 18h. CE - cassette exon, RI – retained intron, A3SS or A5SS - alternative 3′ or 5′ splice site, MXE - mutually exclusive events. **(C)** Comparison of genes with RTI ≥ 0.2 (RTI) and genes with significant changes in RI and CE at the level of nuclear RNA in IM 18h. Only single intersections with RTI are presented. **(D)** Comparison of genes with significant changes in RI (left panel) or CE (right panel) in IM or JTE 18h. **(E)** Sashimi plot for *EIF4H* (left) and quantification of isoform usage (right) in CTRL versus JTE or IM 18h. Isoforms 1 and 2 represent the skipping and inclusion, respectively, of exon 5 (beige highlight). Percent transcripts belonging to each isoform in 3 replicates (mean ± S.E.M.). Significance based on two-tailed Student’s t-test (*, p<0.05; ***, p<0.001).

To determine if alternative mRNA isoforms induced by imatinib are linked to transcriptional readthrough, we performed two types of analyses. First, we asked if retained introns (RI) and cassette exons (CE) detected in nuclear RNA were correlated with genes with RTI≥0.2. Only 95/648 genes had both changes in CE inclusion and high RTI, and only 24/167 genes had both RI and high RTI (Fig. 4C). Interestingly, this set of RI genes includes erythropoietic genes, like UROD, and other genes that will be highlighted below. Intron retention in *DDX17* mRNA is displayed in detail and validated by RT-PCR in Figs. S5C-D. Increased intron retention in *EIF4A2* is likely driven by the presence of five snoRNAs in different introns [43]. Second, we compared sets of alternative mRNA isoforms detected in IM and JTE 18h (see Fig. 4B) to determine the extent of gene overlap. For intron retention (Fig. 4D, left panel), we find that only 4 genes are common between IM and JTE, while 45 intron-retained isoforms are unique to IM. This suggests that transcriptional readthrough does not directly cause intron retention in these genes. Similarly, 324 cases of cassette exon usage were not present in the JTE dataset (Fig. 4D, right panel). However, a substantial overall number (93) of cases of alternative CE usage were detected in the JTE dataset, indicating that a direct inhibition of mRNA 3’ end cleavage may lead to alternative splicing changes. To explore a functionally important example, the mRNA encoding the eukaryotic translation initiation factor 4H (*EIF4H*) harbors a cassette exon (exon 5) that defines two different protein isoforms (Fig. 4E, left), one of which lacks a protein-protein interaction domain required for proper translation initiation [44]. Quantification of splicing events (Fig. 4E, right) and RT-PCR validation (Fig. S5E) revealed that both IM and JTE significantly increase the proportion of isoform 1 and decrease the proportion of isoform 2, containing exon 5. Because 27 of the CE events induced by JTE overlap with those induced by IM, including *EIF4H*, it is therefore possible that these alternative splicing events are directly linked to transcriptional readthrough in both cases. A summary of the observed differential splicing changes attributable to the erythroid gene expression program reveals extensive alternative splicing regulation of numerous up- and down-regulated genes (Fig. S5F).

Given our early detection of imatinib-induced transitions to cell fates (e.g. erythropoiesis), we considered the possibility that some of the observed alternative splicing could be related to the development of imatinib resistance. We analyzed an mRNA-seq dataset from K562 cells treated with a progressively increasing concentration of imatinib over a period of 10 months [45]. These cells are imatinib-resistant and display a large number of alternative splicing changes: 97 mutually exclusive exons, 808 cassette exons, and 162 retained introns (Fig. 5A). Interestingly, none of the identities of retained introns overlap with our IM 18h dataset, suggesting no early steps towards resistance at 18h; however, a large number of cassette and mutually exclusive exons did overlap. This includes *NFE2L1*, which encodes an endoplasmic reticulum-localized protein that translocates to the nucleus upon stress, where it is involved in the regulation of erythroid differentiation as a transcription factor (*NRF1*) (for review see [46]). Results of RT-PCR across the exon validate the change in isoforms (Fig. S5G-H). Gene Ontology analysis revealed many changes consistent with progression to other cell fates, such as transcription factors that promote myeloid differentiation and factors that control proper chromosome segregation (Fig. 5B) for these overlapping alternatively spliced genes. Indeed, many of the overlapping genes are related to erythroid differentiation, showing that development of imatinib resistance involves at least a partial shift to that gene expression program (Fig. 5C). To determine if these changes are comparable to a patient treated with imatinib, we performed mRNA-seq on CD34+ cells isolated at diagnosis and after 6 months of treatment (Fig. 5D). Comparing the different categories of numerous alternative splicing changes present in the patient samples after imatinib treatment, 26 were common with our IM 18h sample, of which 6 genes belonged to the erythroid gene expression program. One example is the translation initiation factor *EIF4A2*, which contains a “poison cassette” exon that harbors premature stop codons, triggering nonsense-mediated decay [47]. Elevated inclusion of this exon is visible in the nascent RNA long-read sequencing dataset and validated by RT-PCR (Figs. 5E, S5I). The relative abundances of the NMD and expressed isoforms are negatively correlated (Fig. 5F), indicating that potentially more EIF4A2 protein is expressed in response to imatinib treatment.

**Figure 5.**
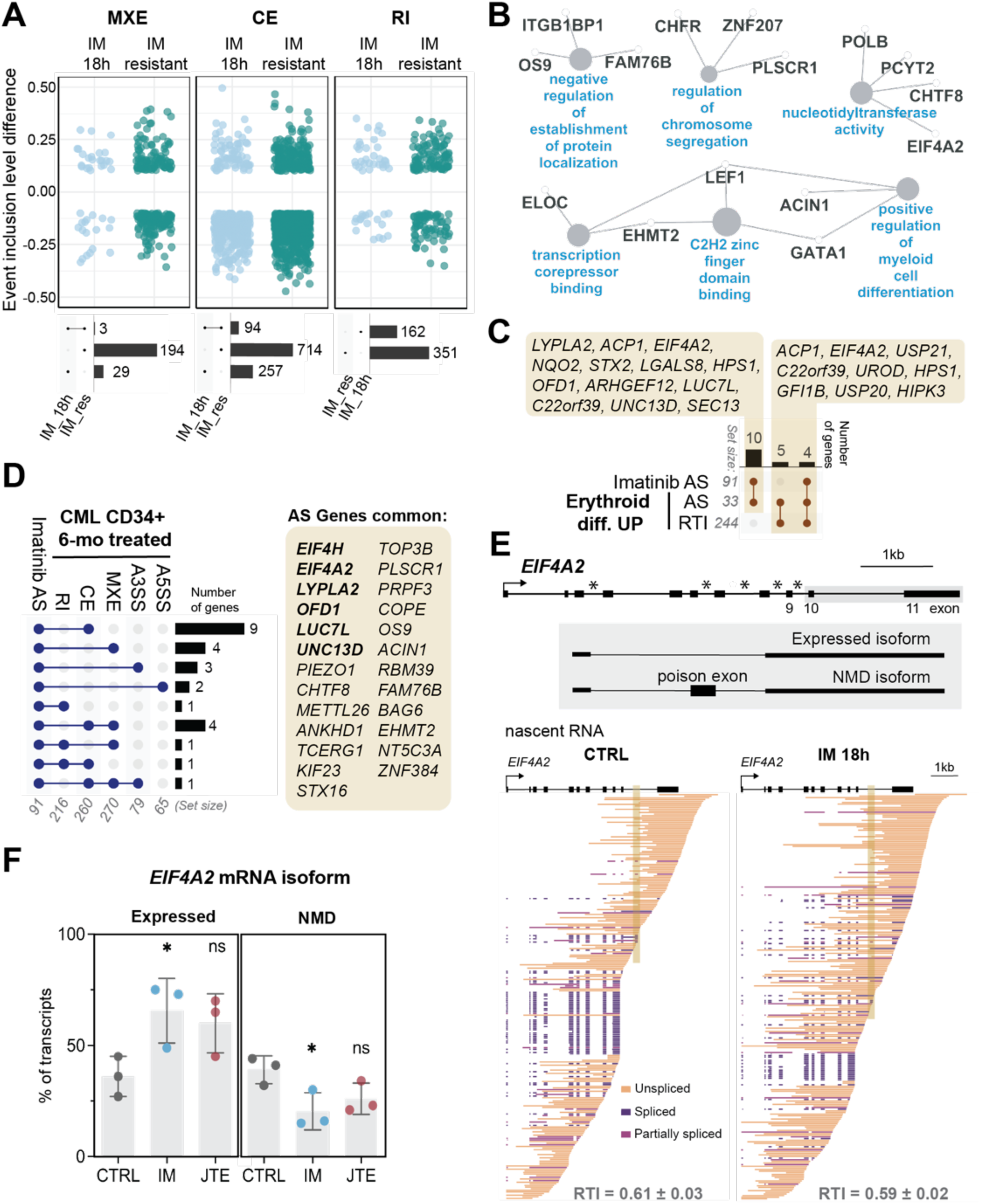
Early changes in alternative splicing control response to Imatinib in a longer time course. **(A)** Comparison of IM-resistant K562 cells (GSE267522) and K562 cells treated with IM for 18h (IM18h). (**Upper panel**) Significant changes in alternative splicing (AS). MXE – mutually exclusive events, CE – cassette exon, RI – intron retention. (**Lower panel**) Upset plots comparing events induced by each condition. **(B)** Enriched Gene Ontology Biological Process terms (blue) for genes that are differentially spliced in both IM-resistant and IM18h treated cells (MXE and CE from A, Lower panel). **(C)** Comparison of genes that are upregulated upon erythroid differentiation (Erythroid diff. UP) and have RTI values ≥ 0.2 (RTImore02) or have significant AS changes upon IM 18h treatment (AS) with genes that have significant changes in CE or MXE for both IM-resistant cells and IM18h (Imatinib AS). **(D)** Comparison of significant AS changes for both IM-resistant and IM18h K562 cells (Imatinib AS) and changes for CD34+-enriched cells from a CML patient at diagnosis and after six months of IM treatment. (**E**) (**Upper panel**) Architecture of eukaryotic translation initiation factor 4A2 (*EIF4A2*). SnoRNAs indicated by black boxes with stars. (**Lower panel**) Downsampled nascent RNA long reads colored according to splicing status, with shaded box over poison exon. RTI values (mean ± S.E.M) for EIF4A2 from three independent experiments are shown. **(F)** Quantification of isoform usage for *EIF4A2*. The NMD and expressed isoforms represent inclusion and skipping, respectively, of the poison exon. Percent transcripts belonging to each isoform in 3 replicates (mean ± S.E.M.). Significance based on two-tailed Student’s t-test (*, p<0.05).

### Imatinib modulates the formation of chimeric mRNAs

Transcriptional readthrough can lead to the formation of RNA chimeras and even produce fusion proteins from those spliced mRNA products [18, 19]. We used bioinformatic tools to search for these “readthrough chimeras” from K562 total and nuclear RNA sequencing datasets as well as CD34+ cells and sought to validate them with our long-read data. Remarkably, the dataset with the most chimeric reads was imatinib-resistant K562 cells and CD34+ from CML patients (Fig. S8A,B). A striking number of chimeras were also detected by short- and long-reads in K562 cells at 18 hours, including the *RBM14-RBM4* chimera in which the 5’SS at the end of *RBM14* exon 1 is spliced to the 3’ splice site of the RBM4 exon 2 (Fig. 6A, 6B, S8A,B). These chimeras correlated with elevated RTI values from the upstream gene (Fig. S8C), and we see that transcription of the downstream gene may increase and/or be coordinated with the upstream gene over a time course of imatinib treatment (Fig. S8D,E).

**Figure 6.**
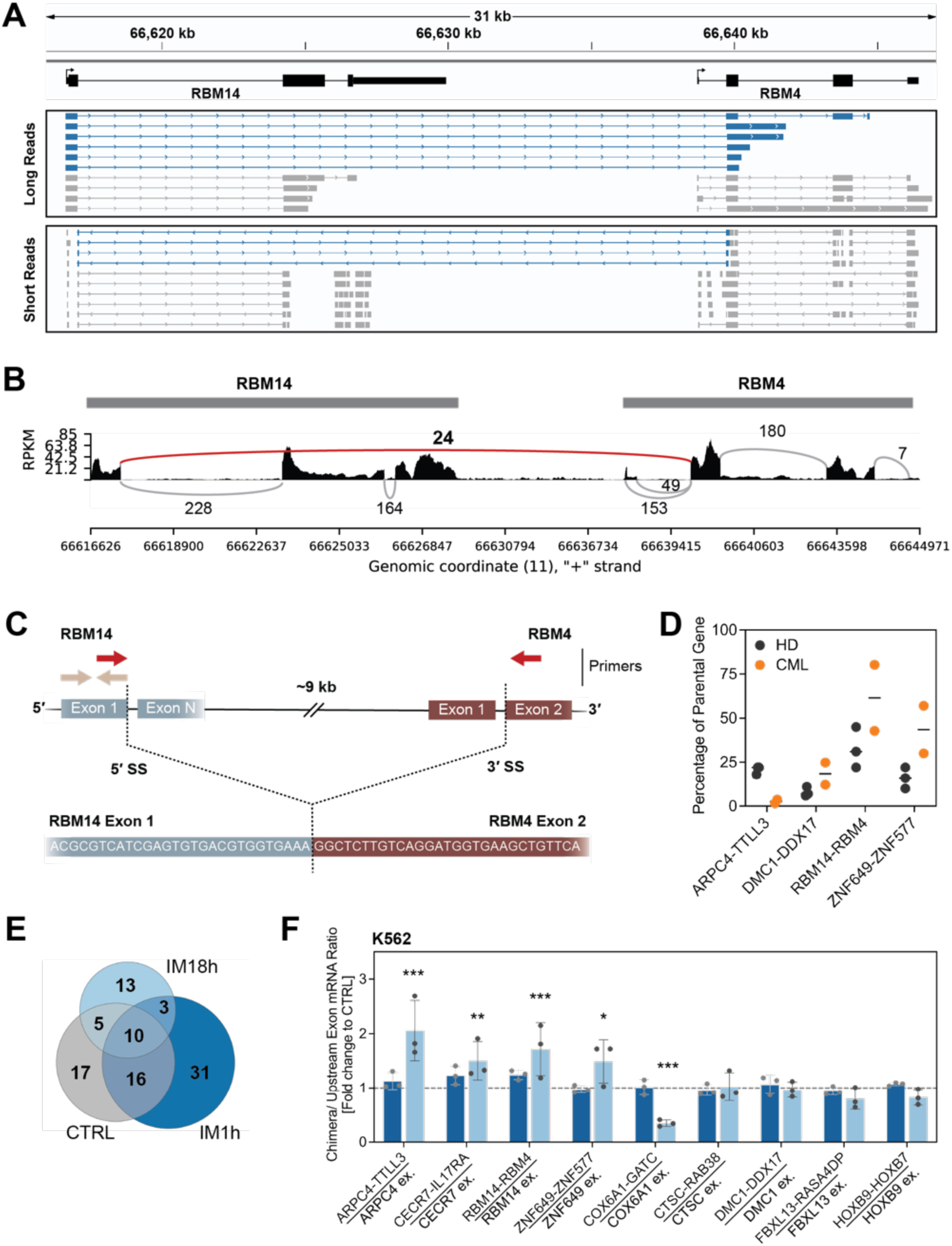
Imatinib-induced formation of readthrough chimeras. **(A)** Examples of reads aligned to RBM14 and RBM4 from long- and short-read sequencing of IM-treated cells. Chimeric reads marked in blue. **(B)** Sashimi plot for RBM14 and RBM4 (chromosome 11: 66616624-66644972) based on a single representative CTRL replicate. Only splicing events detected more than 5 times are shown. Black represents read coverage density in RPKM, gray lines represent splicing events within gene bodies, and the red line represents chimeric splicing. **(C)** Scheme of primer design for real-time PCR semi-quantification (qPCR) of chimera expression. **(D)** Expression of selected chimeras (red primers in C) in CD34+-enriched cells from three healthy donors (HD) and two CML patients (CML), normalized to spike-in controls and presented relative to the expression level of the last exon in the upstream gene (beige primers in C). **(E)** Comparison of readthrough chimeras identified by SOAPfuse analysis of mRNA sequencing for CTRL or IM-treated K562 cells. **(F)** Changes in chimera transcript expression normalized to spike-in control referenced to the last exon in the upstream gene and expressed as fold change upon IM treatment. Level in CTRL as 1.0, marked with the dashed line. Significance based on two-way Anova test (*, p<0.05; **, p<0.005; ***, p<0.001).

To quantify expression and the relative abundance of the single RNA and chimeric RNA isoforms in physiologically relevant cells, total RNA from CD34+ cells from three healthy and two CML patient donors was prepared. Specific primers were designed to amplify either the first exon (or a region of the upstream parental gene) to which the abundance of the chimera – detected with primers across the exon-exon junction – would be normalized (Fig. 6C, Table S4), using reverse transcription followed by quantitative PCR (RT-qPCR). Analysis of four chimeras showed significant expression of *RBM14-RBM4* and *ZNF649-ZNF577* chimeric mRNAs in healthy cells that increased substantially in CD34+ cells from CML patients. In contrast, variable amounts of *ARPC4-TTL3* and *DMC1-DDX17* chimeras were detected in these primary cells. Analysis of our K562 datasets revealed that chimeras are expressed in CTRL (48), IM 1h (60), and IM 18h (31) with significant overlaps in detection (Fig. 6E). Quantification of chimeras by RT-qPCR, using the above method, revealed that some – namely, *ARPC4-TTL3*, *CECR7-IL17RA*, *RBM14-RBM4*, and *ZNF649-ZNF577* – increase significantly by 18 hours of imatinib treatment with little or no increase at 1h (Figs. 6F, S9A), placing them in the category of alternative splicing changes seen at the later timepoint. In contrast, *COX6A1-GATC* is significantly downregulated, while other chimeras do not change. Interestingly, treatment of healthy and CML donor cells with IM 18h did not markedly change absolute levels of chimeras relative to parental genes (Fig. S9A). However, *ARPC4-TTL3* and *RBM14-RBM4* did increase by up to 3-fold relative to CTRL when healthy donor cells were treated for 18 hours and *DMC1-DDX17* increased when CML donors received 1h of imatinib, suggesting the potential for physiological relevance for these cells. We conclude that expression of readthrough chimeras is a specific response to imatinib treatment in K562 and CD34+ cells that is enabled by both transcriptional readthrough and alternative splicing changes.

## DISCUSSION

Here, we have used precision RNA sequencing methods to analyze the earliest changes in gene expression that take place in K562 and CML patient cells upon chemotherapy. Overall, initial responses to imatinib, which directly targets the activity of the *BCR-ABL1* oncogene, are multi-faceted within the first 24 hours. We depict the relative timing and importance of this cascade of gene expression changes as a series of steps and cellular options (Fig. 7). First, long-read sequencing of nascent RNA identified immediate activation of transcriptional readthrough, a blockage or delay in the cleavage of nascent RNA 3’ ends that can be stimulated by cellular stress responses [9, 10]. To our knowledge, this is the first identifiable change in gene expression caused by imatinib, which is known to cause cell death, erythroid differentiation, and chemotherapy resistance at much later time points. Second, we could detect RNA signatures of longer-term imatinib treatment, such as elevation in erythroid gene expression, as early as 1 hour. Previous studies identified development of imatinib resistance in K562 cells [39] involving protection from cell death by ferroptosis [48] as a simultaneous occurrence with erythropoiesis. These unexpectedly early changes in transcriptional readthrough and gene expression may have prognostic value because higher expression of the CD235a erythroid marker is consistent with high efficacy of imatinib treatment and the acquisition of major molecular remission [49]. Third, changes in alternative splicing at 18h of imatinib treatment were also detected in imatinib-resistant K562 cells (10 months) as well as samples from a CML patient collected at diagnosis and after imatinib treatment (6 months), including examples of poison cassette exon usage that reduced the expression of key regulators of translation. Fourth, we discovered chimeric RNAs that were induced by imatinib and elevated in CML patients versus healthy donors, emphasizing the outcome of two coordinated events: ***(i)*** repression of splicing in the upstream gene and ***(ii)*** transcriptional readthrough into the downstream gene; these two events enable the elevated expression of chimeric RNAs and potentially novel protein products in CML cells. We discuss these features in more depth below.

**Figure 7.**
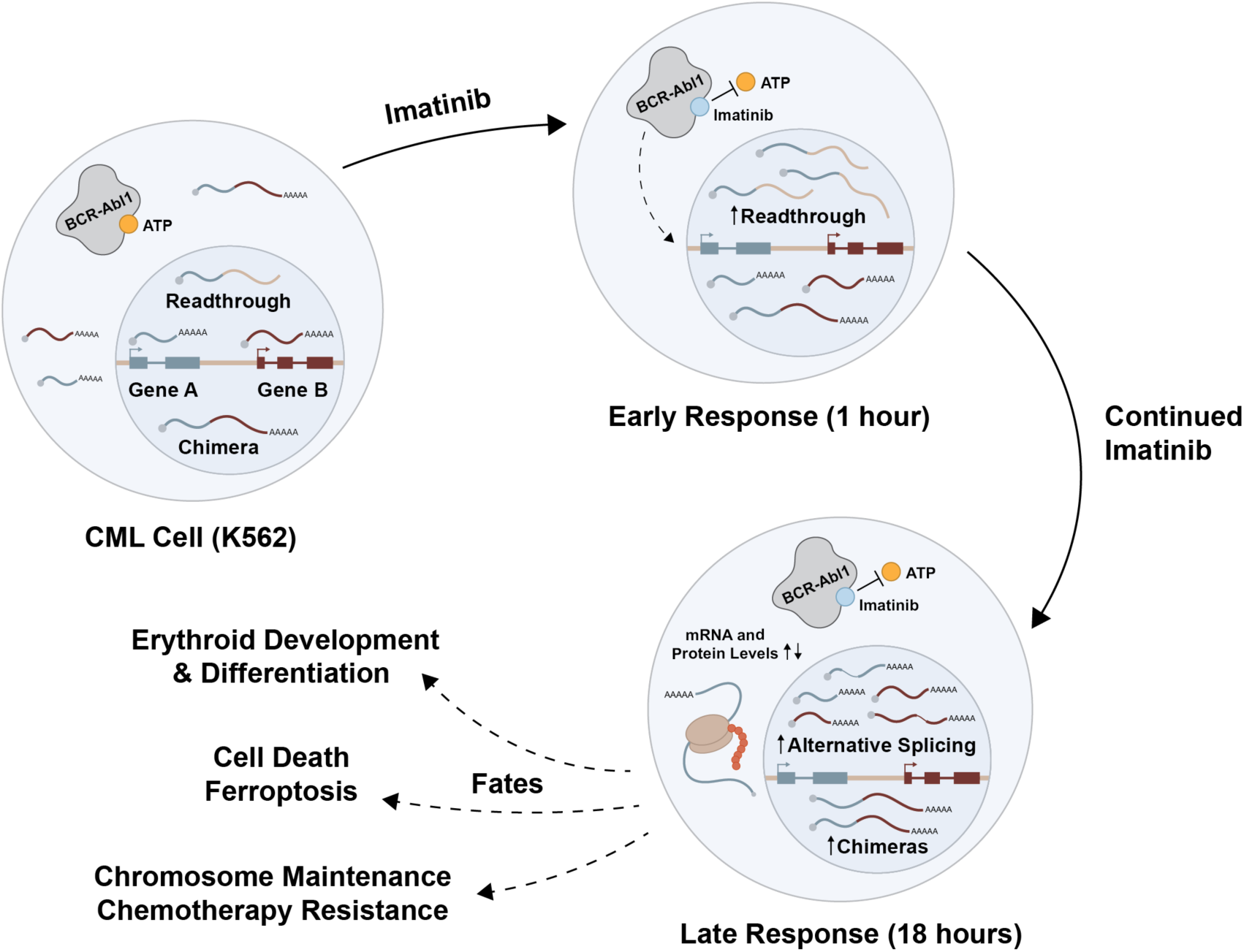
Working model: a cascade of molecular mechanisms triggered by imatinib treatment of CML cells. Transcriptional readthrough is the first major change in gene expression at 1h. At 18 h, intron retention, alternative splicing, and readthrough chimera formation are detected and related to longer-term fates, such as erythropoiesis and resistance to therapy.

We quantified transcriptional readthrough from long-read sequencing of nascent RNA, revealing the proportion of transcripts that are uncleaved at the last annotated polyA cleavage site. This differs from previous determinations of transcriptional readthrough, which quantify short-read density downstream of gene ends [13, 14, 16, 30, 50] because the molecular integrity of each RNA molecule scored is established by the long reads. Our ReadThrough Index (RTI) provides a metric that enabled us to compare the degree of transcriptional readthrough with gene expression, splicing efficiencies, and alternative splicing changes. The observed transcriptional readthrough in cycling K562 cells may be due to the origin of these cells from a human patient in blast crisis, signifying possible cancer stress [36]. Indeed, early observations of significant transcriptional readthrough past annotated gene ends were associated with clear cell renal cell carcinoma [19] as well as stressful treatments, such as heat or hyperosmotic shock [11, 50]. More recently, transcriptional readthrough has been broadly detected in all major cancers, including breast, colon, and liver, where increasing expression of downstream reads is associated with higher morbidity [16]. However, intergenic transcriptional activity is common in healthy tissues as well as virally infected or aging cells [15, 17, 51–53]. Interestingly, when existing readthrough transcripts decrease in cancer, prognoses worsen [16], suggesting there can be positive effects of transcriptional readthrough. One mechanism for phenotypic changes is that transcriptional readthrough into downstream genes can either disrupt the splicing of the downstream gene [8, 14] or lead to readthrough chimeras [18, 19, 53], discussed below.

One outcome of our time-resolved study is that transcriptional readthrough may be mechanistically related to splicing regulation. For example, transcriptional readthrough was associated with inefficient splicing in some of the induced erythroid genes (e.g. *UROD*, *HBE1,* and *FTH1*). Intron retention or inclusion of poison exons reduces mRNA levels by activating nonsense-mediated decay [47], consistent with our data. Transcriptional readthrough correlated with other forms of alternative splicing; DDX17 and EIF4H underwent alternative splicing upon both JTE and imatinib treatments, suggesting that readthrough alone– as opposed to a cellular stress response activated by imatinib treatment– drives alternative splicing in some cases. Long-term responses to imatinib included changes in cassette exon inclusion and intron retention, which are known to [54] the cellular response to therapy [54].

From a pool of alternative isoform changes induced by IM 18h, we identified encoded factors that define erythroid differentiation, such as *GATA1* and *NFE2L1/NRF1* (transcription factors) and *UROD* (a cytoplasmic enzyme involved in the heme biosynthesis pathway). Imatinib caused increased expression of the longer *NFE2L1* isoform, also known as *TCF11*. The Nhe4l domain encoded by the alternative exon alters the spectrum of genes regulated by NRF1/TCF11 and inhibits the development of tumors in mice [54]. Increased expression of *UROD* was observed in *MYCN*-expressing acute lymphoblastic leukemia, and imbalanced synthesis of porphyrins determined cancer cell survival [55]. As postulated recently [56], transcriptional readthrough might facilitate alternative splicing decisions in such cases. We show that alternative mRNA isoforms expressed in imatinib-resistant K562 cells (10 months) occurred much earlier, at 18h of treatment, demonstrating these RNAs as potential molecular signatures of chemotherapy resistance. Interestingly, we identified reduced poison exon inclusion in *EIF4A2* transcripts already within 18h of imatinib treatment. Inclusion of the poison cassette exon between exons 10 and 11 produced a truncated protein that was rapidly degraded in differentiating cardiomyoblasts [57]. Therefore, we infer that reduced inclusion of the *EIF4A2* poison exon would greatly increase protein expression. Given that the promotion of cell growth and proliferation by Bcr-Abl1 involves stimulation of translation [47], this could serve as a rescue mechanism that promotes the progression of cancer despite treatment. This agrees with an earlier observation that alternative splicing of *EIF4A2* was essential for increased protein levels in the development of AML [58].

Effects of transcriptional readthrough and alternative splicing encompass the generation of chimeric transcripts, which has been postulated before [8, 14, 18, 19]. Chimeric transcripts may be hallmarks of RNA processing in oncogenesis and response to therapy. On the other hand, some chimeras were detected in primary CD34+ cells from healthy individuals. Thus, the generation of chimeric transcripts might be a physiological process, with increased pro-oncogenic relevance when other RNA processing steps and protein synthesis surveillance are affected. Increased intron retention accompanying transcriptional readthrough may also contribute because readthrough chimeras require that a 5’ splice site in the upstream gene be spliced to a 3’ splice site in the downstream gene [18]. Thus, the modification of alternative splicing patterns accompanying transcriptional readthrough upon imatinib would drive the generation of new and/or elevated chimeric transcripts. If translated, the resulting chimeric proteins might display gain or loss of function of the original proteins, have different stability, cellular localization, or modified spectrum of interactors. This has been observed in the case of *RBM14-RBM4* [59]. Apart from protein-protein fusions, we also observed lncRNA-protein chimeric transcripts, like lncRNA *CECR7-IL17RA*. Interestingly, stimulation of *IL17RA* cell surface receptor plays a pro-survival role in leukemia, and targeting of IL-17A cytokine increases imatinib’s effectiveness in Bcr-Abl1-positive acute lymphoblastic leukemia [60]. In addition, so-called ‘ripples’ in Pol II transcriptional activity in intergenic regions are associated with changes in chromatin marks and DNA accessibility that could lead to altered protein-coding gene expression [61]. Epigenetic changes accompanying erythroid differentiation [62] could perhaps support CML resistance [63] by broadly adjusting gene expression.

Taken together, the early modifications in RNA processing we observed upon imatinib treatment – including transcriptional readthrough, intron retention, alternative splicing, and readthrough chimera formation – could be exploited as a potential vulnerability after further study. Fusion proteins synthesized from chimeric transcripts can serve as diagnostic markers or neoepitopes for targeted T cell-mediated therapy [64] or anti-cancer vaccines [65]. Similarly, peptides originating from non-sense mediated decay of translated transcripts with retained introns were found to be presented by MHC I [66]. This underscores the significance of the detection of RNA processing changes as early as 1 hour and progressing through an early timeframe of gene expression changes in response to imatinib. Identified changes in RNA processing and detection of isoforms can deliver robust hallmarks of cell responses to therapeutics, like induction of cell differentiation, possibly allowing for early detection of arising therapy resistance.

## MATERIALS AND METHODS

### Cell treatments and fractionation

K562 cells were cultured in Iscove’s Modified Dulbecco’s Medium (IMDM) supplemented with 10% FBS, 100 Units/ml Penicillin, and 100 ug/ml Streptomycin. Cells were treated with 110 mM KCl, 5 μM JTE, or 1 μM Imatinib for 45 min, 1h, 3h, 6h or 18h, as indicated in the figure legends. Each experiment was performed in triplicate to obtain three biological replicates. After washing in cold PBS, 8×10^7^ cells were subjected to fractionation to obtain chromatin-associated nascent RNA, as described [4, 38]. The only modification was an increase in the number of times the pellet was washed with PBS between fractionation steps from one wash to three washes. Fractionation was performed in 4 technical replicates, with 2×10^7^ cells for each replicate. One of the replicates was used to assay the isolated cytoplasmic, nuclear, and chromatin fractions by western blotting. To this end, cold PBS was added to each of the cell and nuclei lysis supernatants to achieve the same final volume. The corresponding volume of PBS was also added to the chromatin pellet. Then, the samples were sonicated on ice at 15% amplitude for 20 sec, and centrifuged for 10 min at 20,000 x g, 4°C. Aliquots of the same volume from each supernatant were incubated with a NuPAGE sample buffer (Thermo #NP0007) at 95°C for 5 min before subjecting them to NuPAGE electrophoresis in a 4-12% Bis-Tris gel (Thermo #NP0322PK2) and MOPS-SDS buffer (Thermo #NP0001). Following transfer, the nitrocellulose membranes were incubated at 4°C overnight with primary antibodies against: GAPDH (SantaCruz #sc-25778), Pol II (4H8) (SantaCruz #sc-47701), Histone H3 (Novus #NB500-171), and alpha-tubulin (Sigma #T6074).

Primary CD34+ cells were enriched from CML patient samples using magnetic beads with anti-CD34 antibodies (Miltenyi #130-046-702) and separated on columns (Miltenyi #130-042-401), following the manufacturer’s protocol. Enrichment up to approx. 70% of CD45+CD34+ cells was confirmed by flow cytometry analysis of the cell surface markers staining. Primary CD34+ cells from healthy donors were expanded in STEM SF II medium (StemSpan #09655) supplemented with the essential growth factors (StemSpan #02691).

### Western blotting analysis of protein level in whole cell lysates

After treatment, the cell pellets were washed in cold PBS followed by lysis at 95°C in the SDS lysis buffer and analyzed by western blotting, as described elsewhere [67]. The following antibodies from Proteintech were used to detect proteins on the nitrocellulose membranes: anti-KEL (#67393-1-Ig), anti-c-Myc (#10828-1-AP), anti-NDC80 (#66960-1-Ig), anti-ATF4 (#10835-1-AP), anti-HBG1 (#25728-1-AP), anti-HBE1 (#12361-1-AP), anti-FTL (#10727-1-AP).

### Library preparation for long-read sequencing on the Pacific Biosciences platform

Nascent RNA was isolated from a chromatin pellet in Trizol (Thermo #15596026) after incubating at 50°C for 10 min with shaking at 1400 rpm, followed by extraction in chloroform and RNA purification using RNeasy Mini kit (Qiagen #74104). DNA was removed by on-column digestion using the RNase-Free DNase set (Qiagen #79254). RNA samples were subjected to library preparation for PacBio long-read sequencing according to the procedure described before [38]. Briefly, rRNA and poly(A+) RNA were depleted using RiboMinus Eukaryote System v2 (Thermo #A15026) and DynaBeads mRNA DIRECT Micropurification (Thermo #61021) kits, respectively, followed by ligation of an adapter with UMI to the 3’ end of RNA by T4 RNA ligase (NEB #M0351L), then reverse transcription by SMARTer PCR cDNA Synthesis kit (Takara/Clontech #634925), and library amplification using Advantage 2 PCR kit (Takara/Clontech #639137). Libraries were subjected to size selection using AMPure XP beads (Beckman Coulter #A63880), to select for fragments longer than 500nt. The PacBio HiFi library was prepared using SMRTbell Express Template Prep Kit 2.0 from PacBio following the manufacturer’s instructions. Libraries were sequence on PacBio RS II and Revio instruments at Yale Center for Genome Discovery.

### Nuclear and total mRNA preparation for short-read sequencing analysis

RNA from control and treated cells, whole cells (total mRNA selected for poly(A+) RNA during library preparation) or nuclei (nuclear RNA depleted of rRNA using RiboMinus Eukaryote System v2) obtained as an intermediate step of the cell fractionation procedure, was isolated following the procedure described above. All treatments were repeated three times as separate biological experiments. Samples were submitted for library preparation at Yale Center for Genome Analysis core facility and sequencing analysis using Illumina Nova-seq HiSeq of paired-end 150 bp fragments.

### Reverse transcription, real-time PCR, and Sanger sequencing

Reverse transcription of total RNA was performed using the SuperScript III Reverse Transcriptase kit (Thermo #18080085), Universal Spike I RNA (TATAA Biocenter #RS25SI), and anchored oligo(dT_18_) (Merck #D2773). Reverse transcription was carried out 50°C for 60 minutes, followed by inactivation at 70°C for 15 minutes. The final cDNA solution was diluted 1:3 for chimeric junction primers and 1:10 for parental exon primers.

The junction sequence of readthrough chimeras predicted by SOAPfuse informed the design of junction-specific primers. Real-time semi-quantitative PCR (RT-qPCR) was performed using the iTaq Universal SYBR Green Master Mix (Bio-Rad #1725121) on a CFX Connect Real-Time PCR Detection System (Bio-Rad), performing 40 cycles of denaturation at 95°C for 10 sec followed by annealing/extension at 60°C for 10 sec. RT-qPCR amplicons were quantified using the ΔΔCt Method, with universal spike-in RNA as the internal control. RT-qPCR products were electrophoresed and resolved on a 1.75% agarose gel. Bands of interest were excised and purified from the gel using the NucleoSpin Gel and PCR Clean-up kit (Takara Bio #740609) according to the manufacturer’s instructions. For junction sequence validation, pre-mixed primer and cDNA samples were sent to Quintara Bioscience for Sanger sequencing.

### Short-read sequencing analysis

#### Short-read sequencing pre-processing

The SRA Toolkit was used to download FASTQ files from the study by Bai J et al. [45] (control samples: SRR29032856, SRR29032857, SRR29032858; Imatinib-resistant samples: SRR29032862, SRR29032863, SRR29032864). For Illumina datasets generated in this study, Illumina adapters were trimmed using FASTP with the following settings: -l 15 -c --detect_adapter_for_pe --trim_poly_g --dedup --trim_front2 0 -w 15. Quality control reports for all datasets were generated using FASTQC. All Illumina datasets were mapped to the hg38 genome using the STAR (v2.7.7a) aligner with settings: --runThreadN 6 -- runMode alignReads --outFilterMultimapNmax 1 --alignSJoverhangMin 8 --alignIntronMin 21 --alignIntronMax 10000 --outSAMtype BAM SortedByCoordinate. The featureCounts program (Subread/2.0.3) was used to quantify the number of reads mapping to each gene with the settings: -T 6 -p --countReadPairs -t exon -g gene_id.

#### Splicing per intron calculation

The Python tool SPLICE-q was used to quantify coverage over the 5’ and 3’ splice sites of each intron [68]. Splicing per intron (SPI) values were calculated based on the SPLICE-q outputs by dividing the number of spliced counts (sum of split coverage over 5’ and 3’ splice sites) by the total number of counts (sum of unsplit and split coverage over 5’ and 3’ splice sites) mapping to that intron. To identify introns with significant changes in splicing, chi-square tests were performed by comparing the number of spliced reads and the number of unspliced reads for pairs of replicates (i.e., replicate one of untreated versus replicate one of treated, etc.). A single, adjusted p-value for each intron was determined using the Benjamini-Hochberg, or False-Discovery Rate, procedure. To be classified as significant, the adjusted p-value had to be less than 0.01. To be considered for SPI analysis, an intron had to have at least 500 reads pooled across all three replicates for both conditions.

#### Alternative splicing analysis

Changes in alternative splicing events for treated samples relative to the control were detected using rMATS-turbo [69] in a Conda environment with the following settings: python=3.7 rmats.py -t paired --readLength 150 --variable-read-length --novelSS -- nthread 6. To be classified as significant, a change in an alternative splicing event had to have an absolute value of inclusion level difference greater than 0.1 and an adjusted p-value less than or equal to 0.05. A read count cutoff was also implemented to keep only events with at least 20 inclusion and skipping reads across per condition, pooled across all three replicates. Only junction reads were considered in the analysis of alternative splicing events. Rmats2sashimiplot was used to quantify and visualize selected changes in splicing for particular genes.

#### Gene expression analysis with DESeq2

Differential gene expression analysis was conducted using RStudio (v4.3.1) and the R/Bioconductor package DESeq2 (v1.40.2). A variance stabilizing transformation was applied using the vst function (with the blind argument set to FALSE). Genes were classified as differentially expressed between control and treated cells if they exhibited a log_2_(Fold Change) greater than 0.6 (upregulated) or lower than −0.6 (downregulated) with an adjusted P-value less than 0.05.

#### Gene ontology term enrichment analysis

Functional annotation analysis of selected genes was performed using g:GOSt (g:Profiler RRID: SCR_006809, https://biit.cs.ut.ee/gprofiler/gost) with Benjamin-Hochberg False Discovery Rate (FDR) set to 0.05. Annotation enrichment was analyzed using Panther (https://www.pantherdb.org) with Fisher’s Exact test requiring p < 0.05. Analysis of protein network enrichment was performed by searching in the ClueGO (RRID: SCR_005748) with GeneOntology (Nov2024) at CytoScape (v 3.10.3) (RRID: SCR_003032) with Benjamini-Hochberg FDR set to 0.05.

#### Identification of readthrough chimera transcripts with SOAPfuse

RNA-sequencing data were preprocessed using FASTP and mapped to the hg38 genome using the STAR aligner with settings: --runThreadN 16 --runMode alignReads --outFilterMultimapNmax 1 -- alignSJoverhangMin 1 --alignIntronMin 21 --alignIntronMax 200000 --readFilesCommand zcat --outSAMtype BAM SortedByCoordinate --outFileNamePrefix ../results/bam/ --sjdbGTFfile {input.gtf}--genomeDir {input.genome_dir} --readFilesIn {input.R1_P} {input.R2_P} -- chimOutType WithinBAM Junctions --chimMultimapNmax 2 --chimSegmentMin 15 -- chimOutJunctionFormat 1 --chimMultimapScoreRange 3 --chimJunctionOverhangMin 15 --chimScoreJunctionNonGTAG −4 --chimNonchimScoreDropMin 10 --alignMatesGapMax 200000 --chimScoreSeparation 5 --chimScoreMin 0. Resulting bam files were used for alternative splicing analysis by rmats2Sashimiplot.

SOAPfuse (v1.26) was used to detect fusion transcripts from preprocessed paired-end RNA-seq data [70]. Alignment database was build using the hg38 human genome and gene annotation files from ENCODE. Reads pooled from biological replicates of the same condition were aligned to the database using the BWA aligner. SOAPfuse analysis settings required minimum 3 span reads and 3 junction covering reads, and a minimal intra-chromosome distance between genes of 500 nt. For comparative analysis, the output list was filtered to include only fusions with at least a total of 6 supporting reads and removed any fusions with ribosomal, small nucleolar, and small nuclear RNAs. To specifically focus on potential readthrough chimeras in each RNA-seq dataset, only chimeras between adjacent genes sharing the same strand and orientation were considered.

### Long-read sequencing analysis

#### Long-read sequencing pre-processing

Adapter sequences were removed using cutadapt (v3.4) with the following settings: cutadapt --cores=8 --discard-untrimmed --no-indels --revcomp -e 0.15. Long reads were mapped to the GRCh38.p14/hg38 genome using minimap2 (v2.22) with settings -ax splice:hq –secondary=no -t 12 -a -u f –seed 14. SAM outputs from minimap2 were converted to BAM and BED files using SAMtools (v1.21) and BEDTools (v2.30.0), respectively. UMICollapse (v1.0.0) was used for deduplication with the following settings: umicollapse - Xmx60g bam --umi-sep. Polyadenylated reads and non-unique reads (i.e., entries with duplicate read names) were removed using a custom script. Pre-processing steps and custom scripts to filter poly(A)+ reads were consolidated in Snakemake.

#### Genome annotation filtering

The “Convert GTF to BED12” feature of the Galaxy web server (https://usegalaxy.org/) was used to generate a BED12 version of the hg38 genome with entries for all transcripts. For splicing analysis, transcript entries were filtered to keep only the canonical transcript as defined by Ensembl for each gene. For readthrough analysis, the transcript with the distal-most 3’ end was used to prevent false positive readthrough identification.

#### Comparison of gene coverage

Analysis of gene region coverage was performed using deepTools2 [Ramirez 2016] in a Conda environment. First, bamCoverage tool was used with the following options: --binSize 40 --normalizeUsing RPKM --effectiveGenomeSize 2913022398. Coverage from 3 replicates was averaged using bigwigAverage. The resulting bigWig files were used to compare coverage of reads within the gene body and 4 kb before the annotated transcription start site (TSS) or after the annotated transcription end site. A coverage matrix was prepared using computeMatrix scale-regions with the following settings: -R hg38_genes.bed (a reference containing only protein-coding genes was used) -bs 40 -b 4000 -a 4000 -- regionBodyLength 7000 --skipZeros --missingDataAsZero. For plotting, the coverage traces were aligned within the gene body region.

#### Splicing status classification and co-transcriptional splicing efficiency calculation

Splicing status classification and co-transcriptional splicing efficiency (CoSE) calculation were performed as described [4]. The splicing status of each long read was assigned based on the number of spliced introns it contained relative to the total number of introns it overlapped. Reads for which all possible introns were removed were classified as “all spliced”, reads for which all possible introns were not yet removed were classified as “all unspliced”, and reads with a combination of spliced and unspliced introns were classified as “partially spliced”. CoSE values were calculated for all introns from canonical transcripts. CoSE represents the number of spliced reads spanning an intron divided by the total number of reads spanning an intron.

#### Readthrough status classification and readthrough index calculation

Long reads were intersected with the last exon of the representative transcript for each gene using bedtools. For reads overlapping multiple genes, only the overlap with the greatest coverage was used for analysis. For each gene, the readthrough boundary was defined as 100 nucleotides downstream of the PAS. A read was classified as “readthrough” if its 3’ end extended beyond the readthrough boundary or classified as “gene body” if its 3’ end did not extend beyond the readthrough boundary. The readthrough index (RTI) was calculated for each gene by dividing the number of readthrough reads by the total number of reads mapping to the last exon of the gene. To be classified as a readthrough gene, the RTI had to be greater than or equal to 0.2.

#### PAS coverage calculation

Long-read coverage in the region downstream of PASs was quantified using a custom Python script. Coverage was normalized to the position 200 nucleotides upstream of each PAS.

## Data availability

Long and short read sequencing raw data are available through the National Center for Biotechnology Information (NCBI) under GEO accession numbers GSE283849 and GSE283813.

## Code availability

Code for RNA-seq data processing and downstream analyses is publicly available at https://github.com/NeugebauerLab/Cancer-stress.

## Acknowledgements

The authors would like to thank members of the Neugebauer and Steitz labs for helpful discussions. The authors are grateful for support from the National Institutes of Health (R01 GM140735 and R01 HL167071 to KMN). MS was supported by the training grant (1T32GM145469-01A1) and an F31 predoctoral fellowship (1F31ES037182-01) from NIEHS. PPB received additional support from a Polish National Science Center grant (2018/30/M/NZ3/00274) and a Senior Fulbright Fellowship (2022/23). PPB is a Forbeck Scholar (2024/27) and a member of the ‘TRANSLACORE’ Cost Action CA21154. The Yale Cooperative Center for Excellence in Hematology providing CD34+ cells was supported by NIDDK Grant U54DK106857-10S1. Data acquisition at Yale Center for Genomic Analysis was supported by the National Institute of General Medical Sciences of the National Institutes of Health under Award Number 1S10OD030363-01A1. The contents of this article are solely the responsibility of the authors and do not necessarily represent the official views of the NIH.

## Author Contributions

P.P.B., M.S., J.L. and K.M.N. conceptualized the study. P.P.B. and J.L. carried out all wet lab experiments, P.P.B., M.S., and J.L. analyzed the data. The original manuscript was written by P.P.B. and K.M.N. and was subsequently edited by P.P.B., M.S., J.L., and K.M.N. The work was by supervised P.P.B. and K.M.N.

## Declaration of Interests

The authors declare no conflict of interest.

## Notes

### Competing Interest Statement

The authors have declared no competing interest.

### Summary of Updates

To meet the journal's requirements, we publish the version of the manuscript originally submitted for review.

