## Supplemental Figures and Tables for "Transcriptional readthrough precedes alternative splicing programs triggered in CML cells by imatinib"

### Supplementary materials

**This PDF file includes:**

Figs. S1 to S9 with text legends
Tables S1 to S4 with text legends

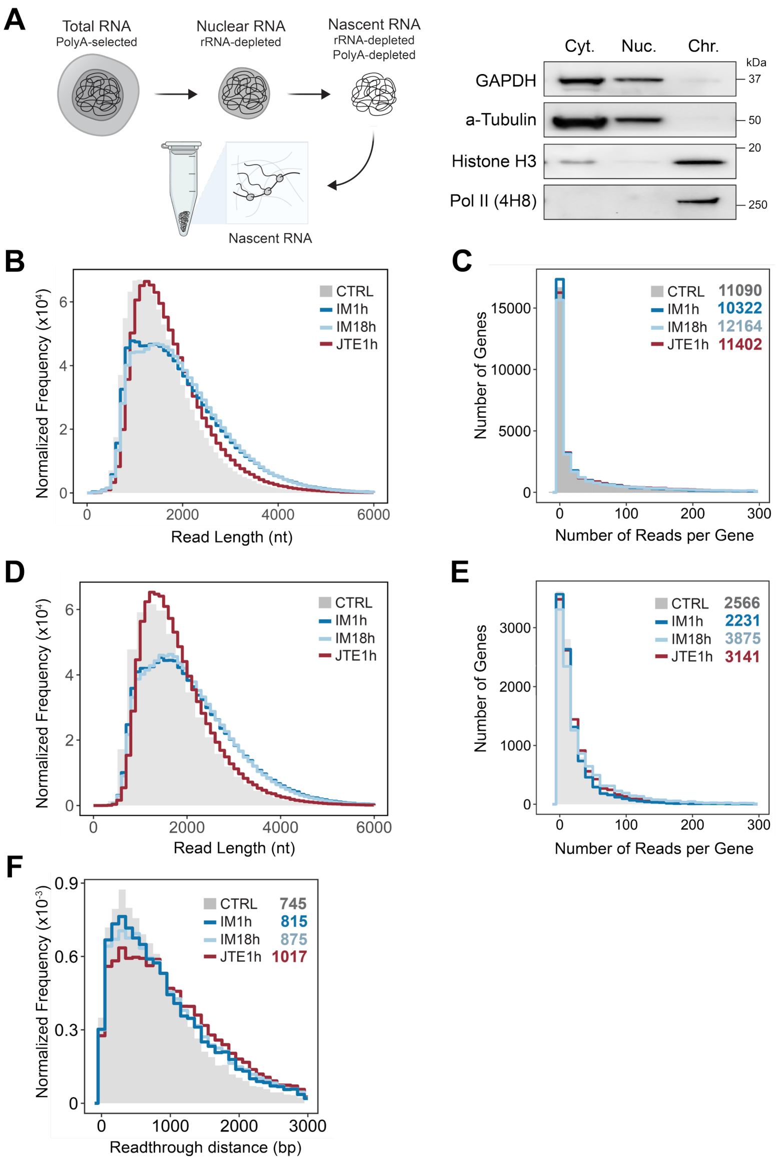


**Fig. S1. Preparation and properties of long-read sequencing datasets.**

**(A)** Representative image of western blotting analysis of protein level in cell fractions (Cyt. – cytoplasm; Nuc. – nucleus; Chr. – chromatin) obtained during isolation of nascent RNA from the chromatin fraction.

Analyses in **(B, C)** include all long reads after poly(A) filtering and splicing intermediate removal, while analyses in **(D-F)** include only filtered long reads that overlap with the last exon of their gene of origin.

**(B, D)** Read length distributions normalized to the number of reads in each condition.

**(C, E)** Read depth distributions. The number of genes with at least 30 reads **(C)** mapping anywhere along the gene or **(E)** mapping to at least the last exon are provided at the top right.

**(F)** Distribution of readthrough distances for expressed genes measured beginning 100 nt after the poly(A) signal of the longest annotated isoform. The median readthrough distance for each condition is provided at the top right.

See Table S1 for numerical data.

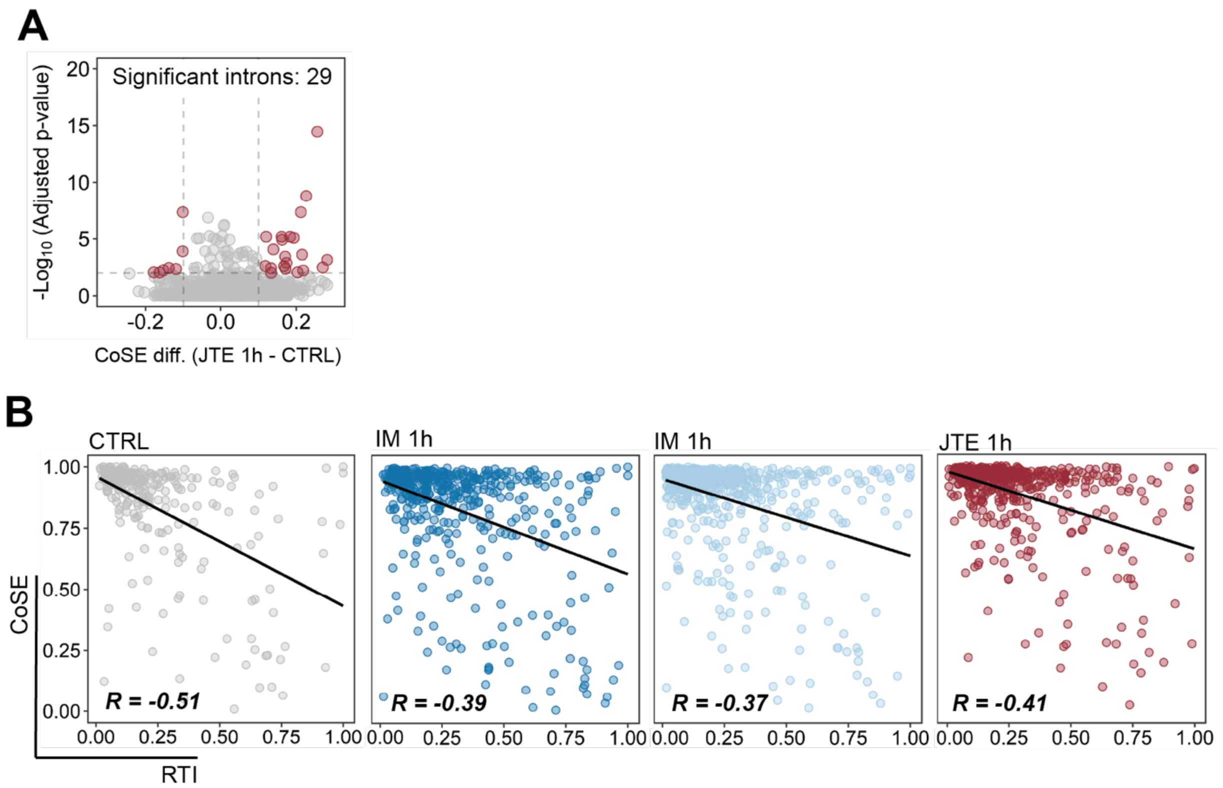


**Fig. S2. Analysis of co-transcriptional splicing in K562 cells before and after treatments.**

**(A)** Change in CoSE value for 1h JTE-treated cells relative to CTRL cells plotted for individual introns. Introns that undergo significant changes in splicing efficiency ($|\text{Change in CoSE}| \geq 0.1$ and adjusted p-value $< 0.05$) are marked in red. Chi-square tests were performed by comparing the number of spliced reads and the number of unspliced reads for pairs of replicates (i.e., replicate one of untreated versus replicate one of treated, etc.). A single, adjusted p-value for each intron was determined using the Benjamini-Hochberg, or False-Discovery Rate, procedure.

**(B)** Scatter plots depicting the relationship between RTI and the average CoSE value for each gene analyzed; R – correlation coefficient value.

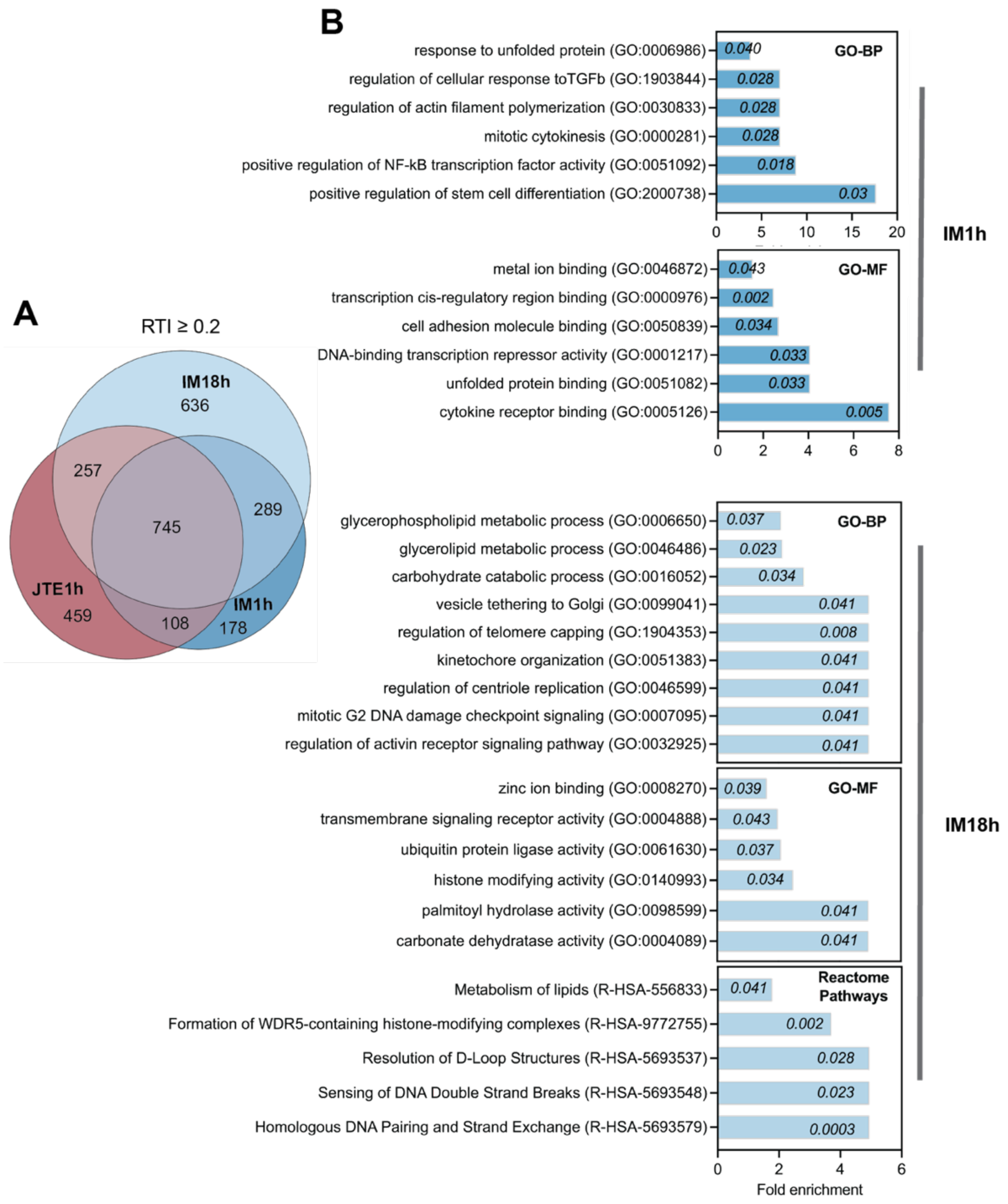


**Fig. S3. Gene Ontology terms enrichment analysis of genes with high transcriptional readthrough after IM 1h and 18h treatments**

**(A)** Comparison of genes with RTI ≥ 0.2 that have at least 30 reads across 3 replicates in each condition. IM – imatinib (1818 of genes in IM18h and 1259 of genes in IM1h), JTE – JTE-607 (1491 of genes).

**(B)** Gene Ontology Biological Processes (GO-BP), Molecular Function (MF), and Reactome Pathways terms enrichment analysis of genes from the intersection of RTI ≥ 0.2 (in **A**) that, excluding the ones that have RTI ≥ 0.2 in CTRL and JTE, are unique to IM 1h (132 genes) or IM 18h (616 genes); Fisher's exact test fold enrichment to all RTI ≥ 0.2 genes identified displayed and FDR value provided in the bars.

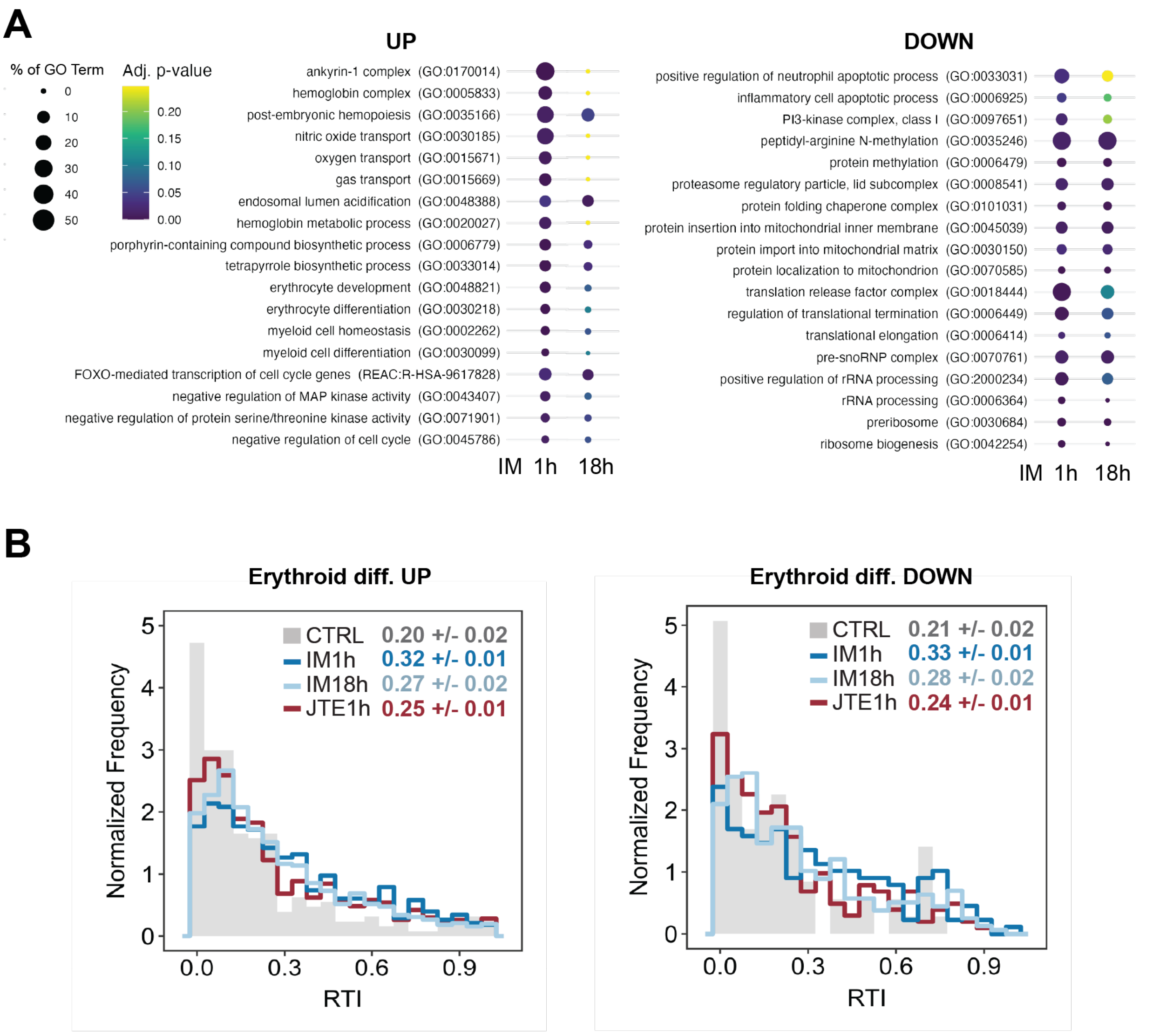


**Fig. S4. Transcriptional readthrough in the context of cell response to treatment.**

**(A)** Gene Ontology Biological Processes terms enrichment analysis of genes with RTI ≥ 0.2 in IM18h or IM1h and RTI difference to CTRL ≥ 0.2 that at the mRNA level in IM18h are upregulated (UP, Log2FC ≥ 0.6) or downregulated (DOWN, Log2FC ≤ -0.6) relative to CTRL. Circle size - percentage of genes representing given term (% of GO Term); Fisher exact test with Bonferroni-Hochberg correction used to estimate significance (Adj. p-value).

**(B)** Transcriptional readthrough index (RTI) value in genes that are differentially expressed upon induction of erythroid differentiation in (An et al. 2014).

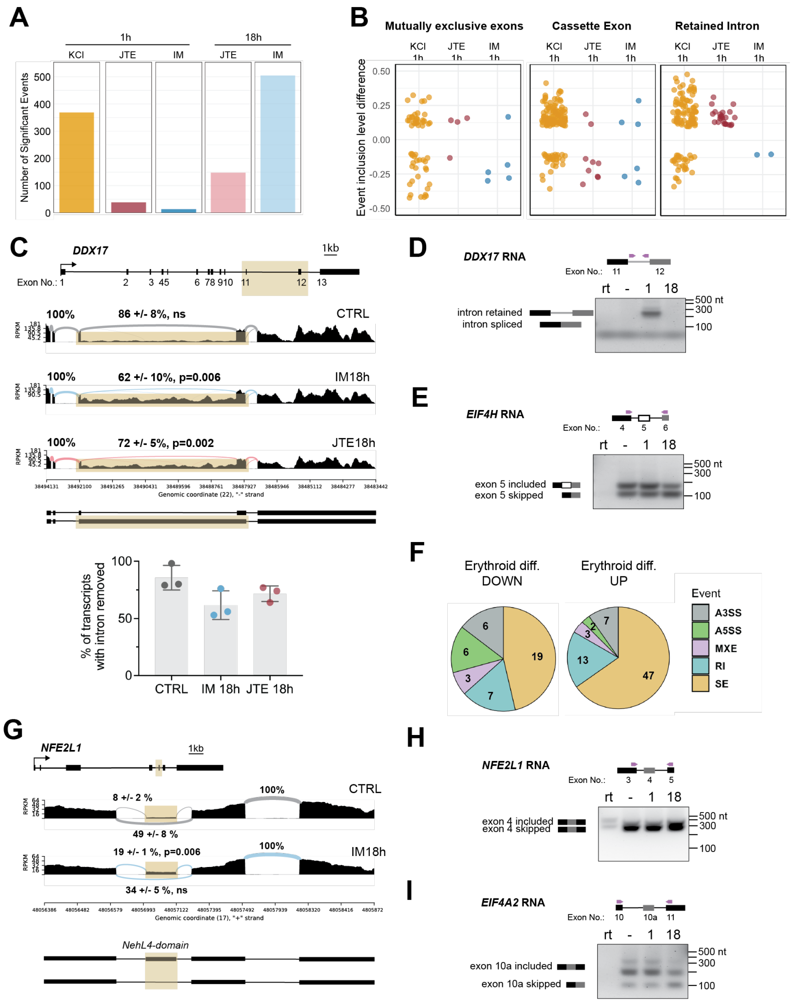

**Fig. S5. Impact of treatment on alternative splicing.**

**(A)** Number of significant (FDR<0.05, 20 reads per event) alternative splicing events in each treatment through analysis of mRNAseq. **(B)** Level difference of exon/intron inclusion (> 0.1) or exclusion (< -0.1); only significant events detected by mRNAseq. **(C)** Sashimi plot showing splicing of DEAD-box RNA helicase 17 (*DDX17*) that is characterized by RTI≥0.2 and intron retention upon IM18h relative to CTRL. Upper panel - coverage and splicing analysis as % of upstream intron on the right; Lower panel – mean value of percent isoforms with intron removed (mRNAseq, n=3). (**D-E**) Images of RT-PCR products from whole cells untreated (-), imatinib-treated cells for 1h or 18h, and no reverse transcriptase control (rt), resolved in an agarose gel for the detection of alternative splicing events by fluorescent staining. Primers (purple) binding indicated above scheme presenting fragment of DDX17 **(D)**, EIF4H **(E)**. **(F)** Alternative splicing events detected in nuclear RNA with significant differences in IM18h relative to CTRL, considering genes identified as UP/DOWN regulated upon CD34+ erythroid differentiation; number of events provided in the piechart. **(G)** Alternative splicing analysis of mRNAseq reads mapping to Endoplasmic reticulum membrane sensor NFE2L1, precursor of transcription factor NRF1 (*NFE2L1*). (**H-I**) Detection of alternative splicing of exon encoding for Nhe4l domain in NFE2L1 (**H**) and poison cassette exon in EIF4A2 **(I)** in by RT-PCR (see above).

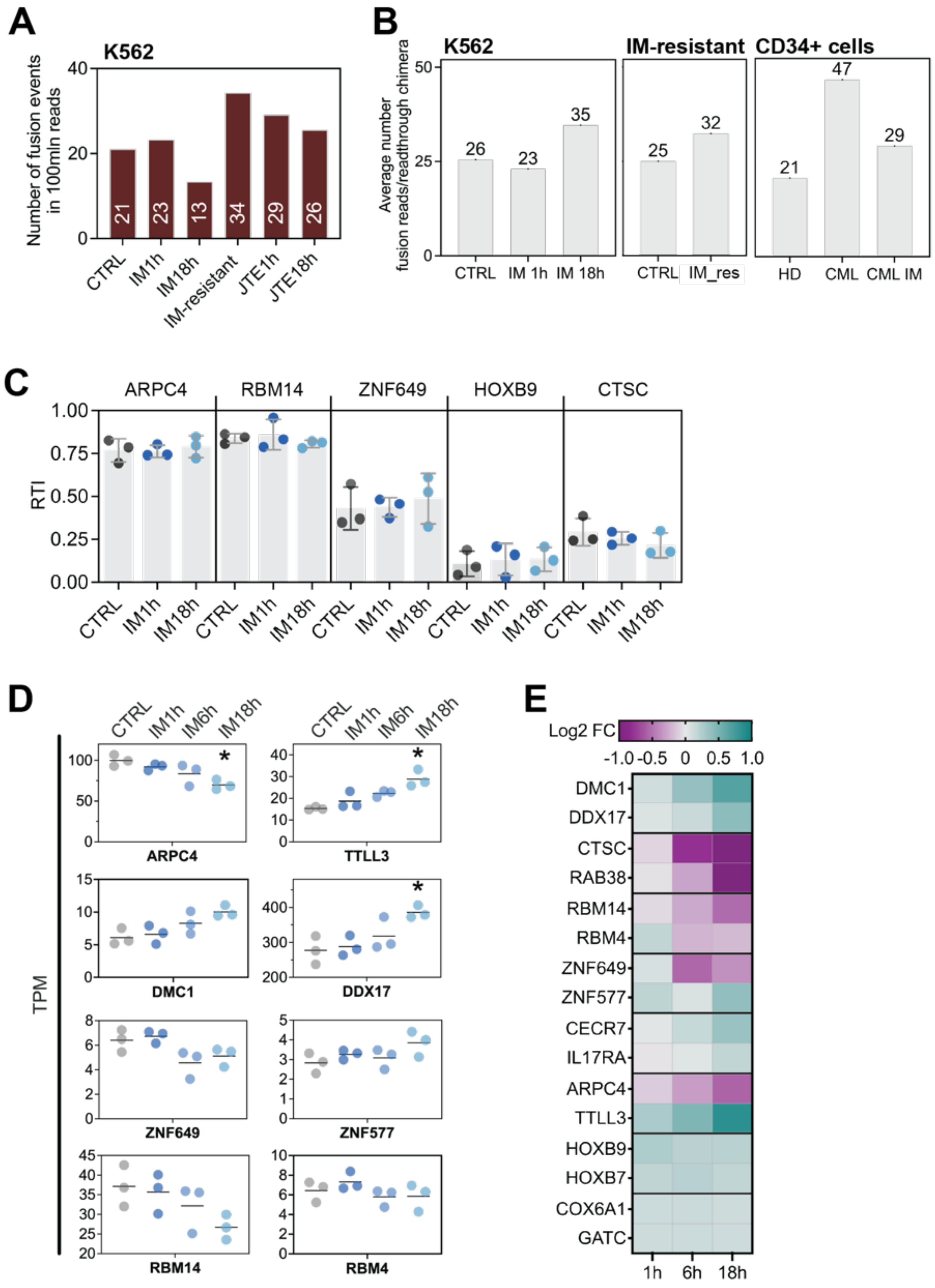


**Fig. S8 Detection of chimeric transcripts (legend next page)**

**Fig. S8. Detection of chimeric transcripts by analysis of short-read mRNA -seq data**

**(A)** Number of readthrough fusion-supporting reads per readthrough chimera in each RNA-seq dataset identified by SoapFuse analysis of short-read sequencing of mRNA from K562 and CD34+ enriched CML cells from patients, untreated (CTRL) or treated with imatinib (IM) or JTE-607 (JTE) for 1h or 18h. Included are also results from analysis of publicly available data from K562 cells imatinib-resistant (Bai J et al., 2024, GSE267522).

**(B)** Average number of reads covering fusion per readthrough chimera. HD – data from CD34+ enriched cells from blood samples of 6 healthy donors (Pellagatti A et al., 2018, GSE114922). CML – our data from sequencing of mRNA from CD34+ enriched cells; CML_IM – 1h imatinib treated.

**(C)** RTI value for selected genes that are upstream in chimera transcripts.

**(D)** Normalized number of reads to the density of sequencing (TPM - transcripts per million reads) for upstream (left) and downstream (right) genes included in the selected chimeric transcripts. Circle - biological replicate; black line – mean value of 3 biological experiments. Student's t-test was used to estimate significance of change upon IM18h to CTRL; * $p<0.05$.

**(E)** Fold change of mRNA level (FC) for chimera forming up- and downstream genes in IM-treated cells for 1, 6 and 18h relative to CTRL.

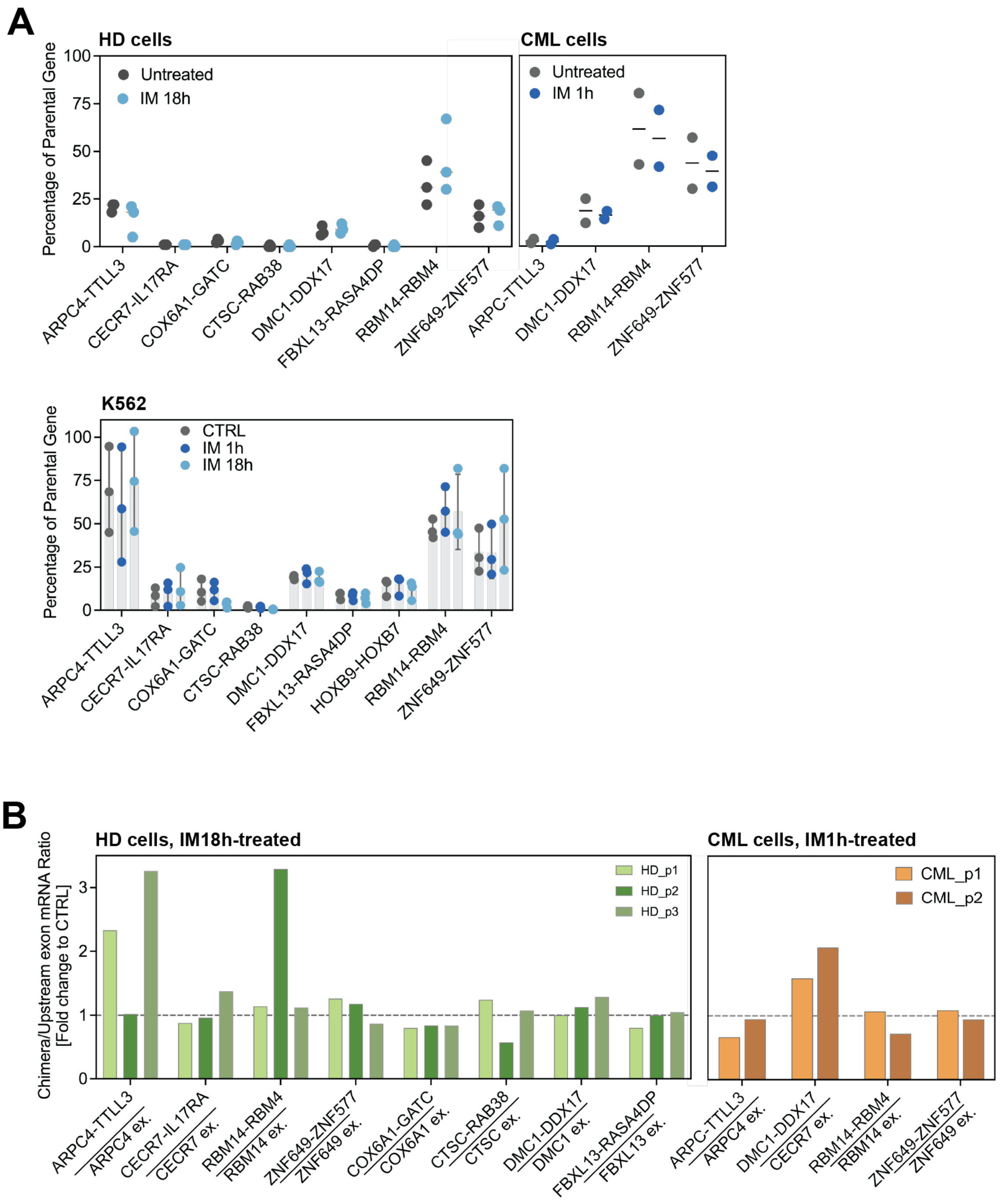


**Fig. S9. Analysis of chimera transcripts level by real time PCR.**

**(A)** Upper panel – primary CD34+ enriched cells from 3 healthy donors (HD) or 2 CML patients, represented by circle, untreated or IM-treated for 18h (in HD) or 1h (in CML). Data normalized to spike-in expressed as percentage of chimera over upstream gene.

**(B)** Comparative analysis of the relative change in chimera abundance upon IM treatment in mRNA from primary cells. Data normalized to spike-in and last exon in the upstream gene presented as fold change to CTRL (set as 1.0 and represented by the dashed line).

**Table S1**

| Sample | raw reads | mapped | w/o poly(A) reads | w/o splicing intermediates |
|---|---|---|---|---|
| CTRL_R1 | 967174 | 416299 | 415587 | 360719 |
| CTRL_R2 | 1109544 | 489147 | 488570 | 410442 |
| CTRL_R3 | 676058 | 284141 | 283657 | 258592 |
| **CTRL_R1-3 sum** | **2752776** | **1189587** | **1187814** | **1029753** |
| KCl 45min_R1 | 1084120 | 469860 | 468992 | 410655 |
| KCl 45min_R2 | 807082 | 343397 | 342671 | 313838 |
| KCl 45min_R3 | 2724768 | 1119386 | 1117365 | 1055596 |
| **KCl_R1-3 sum** | **4615970** | **1932643** | **1929028** | **1780089** |
| JTE 1h_R1 | 1192970 | 519597 | 518931 | 459508 |
| JTE 1h_R2 | 1391536 | 588866 | 587802 | 514227 |
| JTE 1h_R3 | 3202118 | 1298220 | 1295465 | 1147511 |
| **JTE 1h_R1-3 sum** | **5786624** | **2406683** | **2402198** | **2121246** |
| CTRLim_R1 | 296319 | 271399 | 271078 | 229855 |
| CTRLim_R2 | 759545 | 658228 | 657402 | 540074 |
| CTRLim_R3 | 680546 | 599416 | 598488 | 492306 |
| **CTRLim_R1-3 sum** | **1736410** | **1529043** | **1526968** | **1262235** |
| IM 1h_R1 | 907968 | 792025 | 790770 | 673042 |
| IM 1h_R2 | 785490 | 678843 | 677829 | 573238 |
| IM 1h_R3 | 1112676 | 949138 | 947854 | 802732 |
| **IM 1h_R1-3 sum** | **2806134** | **2420006** | **2416453** | **2049012** |
| IM 18h_R1 | 1185161 | 1036352 | 1034219 | 939959 |
| IM 18h_R2 | 1709691 | 1441933 | 1439745 | 1277422 |
| IM 18h_R3 | 1755428 | 1510965 | 1508618 | 1349556 |
| **IM 18h_R1-3 sum** | **4650280** | **3989250** | **3982582** | **3566937** |
| **TOTAL** | **22348194** | **13467212** | **13445043** | **11809272** |

**Table S1. Long-read RNA sequencing datasets and their mapping statistics.** Number of raw reads, mapped reads, poly(A) and splicing intermediates filtered reads from long-read sequencing of nascent RNA (Pacific Biosciences platform) for each described sample. R, biological replicate.

**Table S2**

| | All reads mapped to protein-coding genes | | | | | | | | Gene body reads | | Readthrough reads | |
|---|---|---|---|---|---|---|---|---|---|---|---|---|
| | All | | Spliced | | Unspliced | | Partially spliced | | All | | All | |
| Sample | Median | SD | Median | SD | Median | SD | Median | SD | Median | SD | Median | SD |
| CTRL1 | 1409 | 565 | 1357 | 536 | 1459 | 587 | 1551 | 607 | 1400 | 558 | 1685 | 672 |
| CTRL2 | 1427 | 575 | 1362 | 540 | 1501 | 602 | 1581 | 609 | 1419 | 569 | 1743 | 661 |
| CTRL3 | 1711 | 805 | 1530 | 683 | 1888 | 850 | 1967 | 895 | 1693 | 792 | 2187 | 948 |
| KCL1 | 1390 | 549 | 1352 | 526 | 1427 | 571 | 1551 | 593 | 1383 | 544 | 1678 | 654 |
| KCL2 | 1711 | 840 | 1586 | 758 | 1833 | 892 | 1965 | 915 | 1699 | 831 | 2144 | 1000 |
| KCL3 | 1552 | 726 | 1461 | 657 | 1672 | 782 | 1790 | 807 | 1543 | 718 | 1952 | 893 |
| JTE1 | 1313 | 506 | 1281 | 486 | 1379 | 541 | 1496 | 555 | 1307 | 501 | 1588 | 618 |
| JTE2 | 1813 | 870 | 1650 | 781 | 1939 | 905 | 2034 | 926 | 1784 | 847 | 2320 | 1032 |
| JTE3 | 1694 | 801 | 1573 | 720 | 1837 | 856 | 1939 | 879 | 1669 | 777 | 2243 | 991 |
| IM1h1 | 1871 | 977 | 1613 | 874 | 2068 | 1004 | 2143 | 1024 | 1838 | 961 | 2533 | 1066 |
| IM1h2 | 1798 | 969 | 1542 | 855 | 2009 | 1003 | 2087 | 1014 | 1766 | 953 | 2464 | 1068 |
| IM1h3 | 1836 | 981 | 1574 | 868 | 2044 | 1015 | 2116 | 1023 | 1804 | 965 | 2519 | 1071 |
| IM18h1 | 1915 | 949 | 1703 | 864 | 2131 | 983 | 2211 | 990 | 1891 | 937 | 2512 | 1038 |
| IM18h2 | 1782 | 943 | 1592 | 852 | 1998 | 988 | 2083 | 994 | 1756 | 929 | 2434 | 1041 |
| IM18h3 | 1862 | 946 | 1636 | 851 | 2067 | 981 | 2156 | 979 | 1837 | 935 | 2438 | 1028 |

**Table S2. Read length characteristics of each long-read dataset.** Median with standard deviation (SD) values are shown in nucleotides. The reads described were filtered according to the splicing status or alignment region: Spliced - all introns spliced out, Unspliced - all introns present, Partially spliced - at least one intron spliced out and one intron present, Gene body reads - align within the region of the gene, Readthrough reads - align within the region of the gene covering at least the last exon and extend to the downstream of gene region.

### Table S3

| Sample | Read status | | | |
|---|---|---|---|---|
| | Sequenced | Mapped | Unassigned_ NoFeatures | Unassigned_ Ambiguity |
| Sequencing of mRNA | | | | |
| CTRL_R1 | 22249243 | 19031221 | 1719706 | 1498316 |
| CTRL_R2 | 21683902 | 18333127 | 1875962 | 1474813 |
| CTRL_R3 | 20876632 | 17855236 | 1610102 | 1411294 |
| **CTRL_R1-3 sum** | **64809777** | **55219584** | **5205770** | **4384423** |
| IM 1h_R1 | 22524078 | 18816370 | 2214106 | 1493602 |
| IM 1h_R2 | 25663644 | 21756583 | 2153436 | 1753625 |
| IM 1h_R3 | 25359082 | 21655845 | 2011048 | 1692189 |
| **IM 1h_R1-3 sum** | **73546804** | **62228798** | **6378590** | **4939416** |
| IM 6h_R1 | 26511693 | 22054838 | 2715165 | 1741690 |
| IM 6h_R2 | 22637221 | 19068166 | 2009553 | 1559502 |
| IM 6h_R3 | 20873266 | 17578358 | 1908664 | 1386244 |
| **IM 6h_R1-3 sum** | **70022180** | **58701362** | **6633382** | **4687436** |
| IM 18h_R1 | 21832167 | 18232489 | 2103651 | 1496027 |
| IM 18h_R2 | 22563898 | 18420740 | 2629794 | 1513364 |
| IM 18h_R3 | 21269043 | 17661189 | 2168839 | 1439015 |
| **IM 18h_R1-3 sum** | **65665108** | **54314418** | **6902284** | **4448406** |
| JTE 1h_R1 | 22303801 | 19219839 | 1553035 | 1530927 |
| JTE 1h_R2 | 23519895 | 20323444 | 1543082 | 1653369 |
| JTE 1h_R3 | 22164714 | 19222008 | 1421033 | 1521673 |
| **JTE 1h_R1-3 sum** | **67988410** | **58765291** | **4517150** | **4705969** |
| JTE 6h_R1 | 20133094 | 16971725 | 1846229 | 1315140 |
| JTE 6h_R2 | 22011839 | 18666920 | 1853861 | 1491058 |
| JTE 6h_R3 | 25567472 | 21826935 | 2027655 | 1712882 |
| **JTE 6h_R1-3 sum** | **67712405** | **57465580** | **5727745** | **4519080** |
| JTE 18h_R1 | 21920550 | 18460064 | 2036704 | 1423782 |
| JTE 18h_R2 | 19262757 | 16439848 | 1502147 | 1320762 |
| JTE 18h_R3 | 21378678 | 18095868 | 1884358 | 1398452 |
| **JTE 18h_R1-3 sum** | **62561985** | **52995780** | **5423209** | **4142996** |
| KCl 1h_R1 | 22217976 | 19270518 | 1416170 | 1531288 |
| KCl 1h_R2 | 72943502 | 62953314 | 5081974 | 4908214 |
| KCl 1h_R3 | 49312141 | 42474454 | 3514237 | 3323450 |
| **KCl 1h_R1-3 sum** | **144473619** | **124698286** | **10012381** | **9762952** |
| KCl 3h_R1 | 24077102 | 20822856 | 1573823 | 1680423 |
| KCl 3h_R2 | 26004252 | 22505595 | 1662814 | 1835843 |
| KCl 3h_R3 | 23313941 | 20242385 | 1464866 | 1606690 |
| **KCl 3h_R1-3 sum** | **73395295** | **63570836** | **4701503** | **5122956** |
| **TOTAL** | **1380351166** | **1175919870** | **111004028** | **93427268** |

| Sample | Read status | | | |
|---|---|---|---|---|
| | Sequenced | Mapped | Unassigned_ NoFeatures | Unassigned_ Ambiguity |
| Sequencing of nuclear RNA | | | | |
| CTRL_R1 | 33461963 | 22588484 | 10876630 | 2061588 |
| CTRL_R2 | 32749339 | 22272506 | 10368629 | 2213696 |
| CTRL_R3 | 30988231 | 19803825 | 11211153 | 1967445 |
| **CTRL_R1-3 sum** | **97199533** | **64664815** | **32456412** | **6242729** |
| IM 1h_R1 | 28385612 | 20421694 | 7869067 | 1858137 |
| IM 1h_R2 | 34601349 | 22828792 | 11745409 | 2162499 |
| IM 1h_R3 | 30044816 | 20924712 | 8990724 | 2052763 |
| **IM 1h_R1-3 sum** | **93031777** | **64175198** | **28605200** | **6073399** |
| IM 18h_R1 | 36035776 | 23580149 | 12268138 | 2300791 |
| IM 18h_R2 | 30929803 | 20075848 | 10693243 | 2036782 |
| IM 18h_R3 | 25304023 | 16175394 | 9014972 | 1682168 |
| **IM 18h_R1-3 sum** | **92269602** | **59831391** | **31976353** | **6019741** |
| JTE 1h_R1 | 35284411 | 23709633 | 11556796 | 2157487 |
| JTE 1h_R2 | 30225874 | 20855304 | 9261584 | 1981171 |
| JTE 1h_R3 | 36189754 | 25103205 | 10927819 | 2480259 |
| **JTE 1h_R1-3 sum** | **101700039** | **69668142** | **31746199** | **6618917** |
| KCl 1h_R1 | 28121738 | 19251170 | 8806106 | 1758029 |
| KCl 1h_R2 | 31773769 | 21728391 | 9939890 | 2129214 |
| KCl 1h_R3 | 32496131 | 20518683 | 11971735 | 2040885 |
| **KCl 1h_R1-3 sum** | **92391638** | **61498244** | **30717731** | **5928128** |
| **TOTAL** | **953185178** | **639675580** | **311003790** | **61765828** |

**Table S3.**

**Short-read RNA sequencing and mapping statistics.** Number of raw reads sequenced on the Illumina platform, counts of mapped reads, number after filtering out the reads mapped to a region that is not annotated in hg38 annotation (Unassigned_NoFeatures), Number after filtering for reads mapped to multiple locations in hg38 reference genome (Unassigned_ambiguity). R, biological replicate.

### Table S4

| **Chimeric Junction** | Sequence | Amplicon Size |
|---|---|---|
| ARPC4-TTLL3_F | GAAGCTGTCAGTCAATGCCC | |
| ARPC4-TTLL3_R | TCTGAGCCGGTTCATGTGAG | **111** |
| CECR7-IL17RA_F | GGACCAGCTGACTAAGGAGC | |
| CECR7-IL17RA_R | CTTGTTGGGTGTGGGCAAAG | **177** |
| COX6A1-GATC_F | TGTGTACCTGAAGTCGCACC | |
| COX6A1-GATC_R | CGATAGCTTTCTCCAGTCGC | **184** |
| CTSC-RAB38_F | CAGACCCCAATCCTAAGCCC | |
| CTSC-RAB38_R | ATGCACCCATAGCTTCTCGG | **112** |
| DMC1-DDX17_F | AGCTTGCGAAAGGGAAGAGG | |
| DMC1-DDX17_R | TGTCAGCCTTGCTACTTCCG | **198** |
| FBXL13-RASA4DP_F | GCTGCAAACAACTCCGGATC | |
| FBXL13-RASA4DP_R | TGTGTCTGTGCTACTGTCTTCC | **133** |
| HOXB9-HOXB7_F | CAGTTTGGAAACTTCGGCGG | |
| HOXB9-HOXB7_R | TCTGGTAGCGGGTGTAGGTC | **155** |
| RBM14-RBM4_F | TTTTCGTGGGCAATGTGTCG | |
| RBM14-RBM4_R | TCACATTCCAGCACCTTCCC | **203** |
| ZNF649-ZNF577_F | ACATCACAGAACTCACACGGG | |
| ZNF649-ZNF577_R | AGCCTGGAAGCTCCTTCTTC | **103** |

| **Upstream Parental Exon** | Sequence | Amplicon Size |
|---|---|---|
| ARPC4_Ex5_F | ACTTCCACACAGAGCAGATGT | **130** |
| ARPC4_Ex5_R | TCTTCAGCCACAATGCGGG | |
| CECR7_Ex1_F | AGAGAACACAGCCGAAGTGG | **89** |
| CECR7_Ex1_R | CCAGGAACCAGATGTAGCCC | |
| COX6A1_Ex2_F | AGCTCGCATGTGGAAGACTC | **87** |
| COX6A1_Ex2_R | TGGTGCGACTTCAGGTACAC | |
| CTSC_Ex5_F | CAAACCTGCACCACTGACTG | **113** |
| CTSC_Ex5_R | CTTGGTTTCGAACAGGACTGAC | |
| DMC1_Ex_F | ACCCATTGGGGGACACATTC | **71** |
| DMC1_Ex_R | GCTCTCCTCTTCCCTTTCGC | |
| FBXL13_Ex_F | CAGCAATGGAGATGTTATCGGC | **122** |
| FBXL13_Ex_R | GATCCGGAGTTGTTTGCAGC | |
| HOXB9_Ex1_F | CAGTTTGGAAACTTCGGCGG | **107** |
| HOXB9_Ex2_R | GATCCGGCCTCTCTTTGTCC | |
| RBM14_Ex1_F | GAAACAGTTCGCCTTCGTGC | **160** |
| RBM14_Ex1_R | CGACACATTGCCCACGAAAA | |
| ZNF649_Ex1_F | CCAGTGCCGGGGTTTAAGG | **70** |
| ZNF649_Ex1_R | CTGATTTTCTTCTGGCTACACCT | |

| **Alternative splicing validation** | Sequence | Amplicon size |
|---|---|---|
| EIF4H_Ex4_F | GTGTGGACATTGCAGAAGGC | |
| EIF4H_Ex6_R | GCCCCCTAAGAAGTCATCCC | **101** |
| DDX17_Ex11_F | CTGGAAAGGCACCCATCCTT | |
| DDX17_Ex12_R | CCTCTGAGCTGTTTGGATAGTCA | **105** |
| NFE2L1_Ex3_F | CTGATTGACATCCTTTGGCGAC | |
| NFE2L1_Ex5_R | GGCTCACTCTCACTAGGCAC | **257** |
| EIF4A2_Ex10_F | ATGTGCAACAAGTGTCTTTGGT | |
| EIF4A2_Ex11_R | CTCCCAAATCGACCCCCTCT | **94** |

| **Downstream Parental Exon** | Sequence | Amplicon Size |
|---|---|---|
| TTLL3_Ex_F | ACAATCCAAGGCTGCTACCC | **131** |
| TTLL3_Ex_R | CCCATCGCTGAGCTATCCAG | |
| DDX17_Ex2_F | TTTGGTAATCCTGGGGAGCG | **108** |
| DDX17_Ex2_R | TGTCAGCCTTGCTACTTCCG | |
| RBM4_Ex2_F | TGGGAAGGTGCTGGAATGTG | **185** |
| RBM4_Ex2_R | GACTGATGTTGCCCACATGC | |
| ZNF577_Ex_F | CCAGTGGTAGCTTGACCTGA | **123** |
| ZNF577_Ex_R | GGCCTTGCCCATTTCGTTCA | |

**Table S4.**
**Primers designed for real-time PCR analysis of transcript level and alternative splicing.** F - forward, R - reverse, Ex – exon.
